# Annexin A2 mediates RIG-I-like receptor responses to viral infection

**DOI:** 10.1101/2025.10.28.684968

**Authors:** Georgie Wray-McCann, David Michalík, Alexandra L. McAllan, Linden J. Gearing, Roxanne Hsiang-Chi Liou, Nicole K. Campbell, Sarah Straub, Hani Hosseni-Far, Antony Matthews, Michelle D. Tate, Sophia Davidson, Roman Fisher, Carrie Bernecky, Jan Rehwinkel, Paul J. Hertzog, Natália G. Sampaio

## Abstract

RIG-I and MDA5 are critical sensors of virus infection and detect many important human pathogens including SARS-CoV-2, influenza A virus (IAV), Zika virus and West Nile virus (WNV). MDA5 is activated by binding to double-stranded RNA generated during infection and induces strong pro-inflammatory and antiviral responses mediated largely via type I interferons. Activation of MDA5 is also implicated in monogenic (e.g. Aicardi-Goutières syndrome) and complex polygenic (e.g. systemic lupus erythematosus) autoinflammatory diseases, demonstrating the importance of appropriate regulation of this pathway. Yet, how MDA5 is regulated is poorly defined. We employed SILAC-mass spectrometry to discover MDA5 binding partners in virus-infected cells. Our screen revealed Annexin A2 (ANXA2) as an interactor of MDA5 specifically during infection. Direct binding of ANXA2 and MDA5 was recapitulated *in vitro* and required Ca^2+^, and ANXA2 promoted MDA5 oligomerisation in cells. Loss of ANXA2 in human cells significantly blunted interferon responses to EMCV, WNV and IAV, demonstrating a role for ANXA2 in activation of both MDA5 and RIG-I. Using human induced pluripotent stem cells, we found that ANXA2 was required for potent neuronal cell responses to WNV infection and macrophage responses to multiple viral pathogens. Thus, we find that ANXA2 is a critical mediator of RIG-I-like receptor pathways.

## Introduction

The innate immune system provides early and immediate protection from pathogens. This is largely mediated by pattern recognition receptors (PRRs) that can detect common features of pathogens, which are normally absent in the host. The detection of viral infection by PRRs occurs mainly through the sensing of aberrant or mislocalised nucleic acids, for example unusual RNA species produced during infection or DNA in the cytosol. Activation of these pathways triggers a proinflammatory and antiviral response, centrally involving the release of type I interferons (T1-IFN) and type III interferons. Melanoma differentiation-associated protein 5 (MDA5) is a cytosolic PRR that can detect many viruses (*1*). It belongs to the retinoic acid inducible gene I (RIG-I)-like receptor (RLR) family, along with RIG-I itself and laboratory of genetics and physiology 2 (LGP2) (*2*). MDA5 detects double-stranded RNA (dsRNA) in the cytosol that is generated during virus infection, but is low or absent during homeostasis (*3*). MDA5 can detect infections by many different virus families, including Coronaviridae (e.g. SARS-CoV-2), Flaviviridae (e.g. Zika virus, West Nile virus), and Picornaviridae (e.g. Rhinovirus, Encephalomyocarditis virus, Poliovirus) (*1*). In contrast, RIG-I is activated by 2’-O unmethylated duplex RNA containing tri- or di-phosphates at the 5’ end (*2*). These RNA species are generated during replication of specific RNA viruses (e.g. Influenza A virus) but are absent or masked in host cytosolic RNA. The combination of MDA5 and RIG-I sensing allows comprehensive innate immune detection of most RNA viruses in cells, and many viruses activate at least one of these receptors.

Loss of function of MDA5 leads to repeated life-threatening susceptibility to viral infections, demonstrating the importance of this receptor in our defence against viruses (*4, 5*). On the other hand, although MDA5 is critical to our protection, it is also implicated in several autoimmune and autoinflammatory diseases (*1*). Continuous MDA5 activation leads to excessive and chronic production of T1-IFN, which has damaging effects. Single-nucleotide polymorphisms in the *IFIH1* gene that encodes MDA5 have been associated with type 1 diabetes (*6–8*) and systemic lupus erythematosus (*9*). Furthermore, rare activating mutations of MDA5 lead to severe disorders such as Aicardi-Goutières syndrome and Singleton-Merten syndrome (*10*). Finally, in some virus infections, chronic activation of the innate immune system can lead to a damaging acute response (often referred to as the ‘cytokine storm’), which can be fatal, as has been demonstrated for SARS-CoV-2 (*11, 12*). Thus, understanding the cellular mechanisms employed to regulate RLR activity has implications for a range of human diseases.

It has been reported that MDA5 specifically recognises long dsRNA structures, where the length allows multiple MDA5 monomers to bind and oligomerise to form an activated platform for downstream signalling (*3, 13*). MDA5 contains two caspase activation and recruitment domains (CARDs), followed by a helicase domain and a carboxy-terminal domain. The latter two domains are responsible for dsRNA ligand recognition and binding, whereas the CARDs mediate downstream signalling via interaction with mitochondrial antiviral signalling protein (MAVS). Activated MAVS also oligomerises, recruits additional elements, and ultimately results in the activation of transcription factors like interferon regulatory factors (e.g. IRF3 and IRF7) and NF-κB (*14*). These transcription factors induce the expression of T1-IFNs, proinflammatory cytokines and other antiviral proteins. Upon binding to its RNA ligand, RIG-I also engages MAVS via its own CARDs and activates the same downstream signalling pathway (*2*).

PRRs, due to their capacity to elicit strong and potentially damaging proinflammatory responses, have tightly regulated pathways. Furthermore, they often require accessory proteins for complete pathway activation. Protein regulators of RLRs mostly function by catalysing post-translational modifications, such as phosphorylation or ubiquitination of RLRs, or through direct interactions that promote or inhibit RLR activity (*2*). Regulators of RIG-I have been well characterized, but these are not as well defined for MDA5 (*2*). MDA5 activation requires dephosphorylation at specific sites, mediated by PP1α/γ (*15*). K63-linked ubiquitination and SUMOylation of MDA5, mediated by TRIM65 and TRIM38, respectively, is also required for full activation (*16, 17*). It was also shown that MDA5 ISGylation (the conjugation of the ubiquitin-like protein ISG15), mediated by HERC5 in humans (HERC6 in mice), is required for MDA5 oligomerisation and full activation and response to infection, both *in vitro* and *in vivo* (*18, 19*). The termination of MDA5 signalling is regulated by ubiquitination (*20, 21*). Additionally, IFI27 (*22*) and GLUT4 in muscle cells (*23*) have been reported to inhibit MDA5 via substrate sequestration or segregation away from MAVS, respectively.

An important step in MDA5 activation is its oligomerisation to allow multiple MDA5 CARDs to interact and activate MAVS. 14-3-3η interacts with MDA5 CARDs and promotes its oligomerisation and translocation to MAVS (*24, 25*). PACT also promotes MDA5 oligomerisation by enhancing its binding to dsRNA (*26*). Recently, ADAM9 was found to have a similar function, with loss of ADAM9 resulting in reduced MDA5 oligomerisation, dampened T1-IFN responses, and increased susceptibility to EMCV *in vivo* and *in vitro* (*27*). ZCCHC3 promotes RLR activation by interacting with RNA ligands and promoting liquid phase separation, which co-enriches a subcellular region with RNA ligands and RLRs, and facilitates receptor oligomerisation and interaction with MAVS (*28*). The requirement of RLR movement to a distinct subcellular localisation for its encounter with dsRNA may be particularly important for MDA5 activation, which is believed to require sufficient dsRNA above a certain dose threshold in order to trigger effective signalling from the receptor (*29*). It is noteworthy, however, that several of these studies have been performed in overexpression systems and using artificial dsRNA mimics, whereas investigations using endogenous proteins and live virus infections are less common. This may hinder our discovery of additional critical processes that occur specifically during virus infections, but which are not evident when using reductionist biological systems.

Using a mass spectrometry approach to investigate endogenous MDA5, we identified interactors of MDA5 upon virus infection. Of these, ANXA2, a pleiotropic Ca2^2+^-binding protein, was found to directly interact with MDA5 upon virus infection and promote MDA5 oligomerisation. Loss of ANXA2 compromised the cell’s ability to mount an interferon response to multiple viruses, and hindered both MDA5 and RIG-I responses to RNA ligands. We reveal here that ANXA2, previously not known to play a function in innate immune signalling pathways, is a mediator of RLR activation.

## Results

### ANXA2 interacts with MDA5 upon virus infection

In order to discover regulators of MDA5 signalling, we developed an endogenous immunoprecipitation (IP) approach for MDA5. The human monocytic cell line THP1 was selected as it expresses high levels of MDA5 at baseline and can be infected with EMCV, a prototypical picornavirus that activates MDA5, but not RIG-I (*30*). This method isolated endogenous MDA5 and co-precipitated additional binding partners upon virus infection (Fig 1A,B). We employed SILAC mass spectrometry to quantifiably identify proteins that interacted with MDA5 upon infection. THP1 cells were differentially labelled with either light or heavy isotope, and left uninfected or infected with EMCV at multiplicity of infection (MOI) of 10 for 20 h. Equal concentrations of cell lysate from each condition were combined and used for IP of endogenous MDA5. The MDA5-bound sample was then analysed by mass spectrometry to identify co-precipitating proteins.

**Figure 1:**
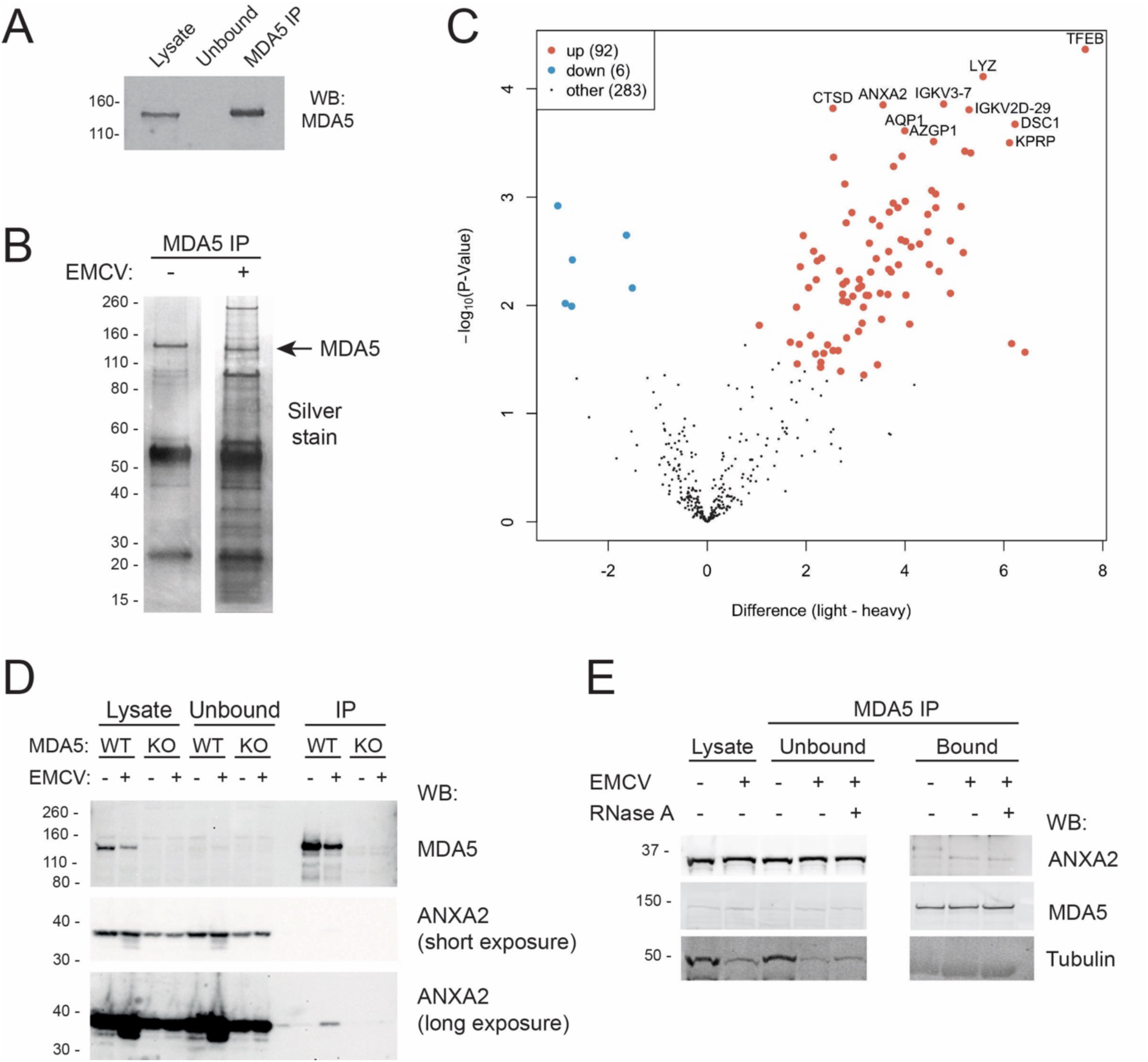
SILAC mass spectrometry identifies ANXA2 as a binding partner of MDA5 upon EMCV infection. (A) Endogenous MDA5 was immunoprecipitated from THP1 cells and eluted with 200 mM glycine, pH 2. Cell lysate, unbound sample and bound sample (IP) were separated by SDS-PAGE and analysed by western blot for MDA5. (B) THP1 cells were left uninfected or infected with EMCV (MOI = 10) for 20 h. MDA5 IP was performed and samples separated by SDS-PAGE and the gel was silver stained to detect co-eluted proteins. (C) Light (R0K0) or heavy (R10K8) SILAC-labelled THP1 cells were either left uninfected or infected with EMCV (MOI = 10) for 20 h, respectively. Equivalent cell lysate from each condition was combined and MDA5 was immunoprecipitated as in (A). The volcano plot shows the differentially enriched proteins in three independent biological replicates as measured by mass spectrometry. (D) Wild-type or MDA5-KO THP1 cells were uninfected or infected with EMCV (MOI = 5, 20 h). MDA5 was immunoprecipitated and cell lysate, unbound sample, and bound sample (IP) were separated by SDS-PAGE and analysed by western blot for MDA5 and ANXA2. (E) THP1 cells were left uninfected or infected with EMCV (MOI = 5, 20 h). Cell lysate was treated with RNase A or mock-treated, and samples separated by SDS-PAGE and analysed by western blot for ANXA2, MDA5, and tubulin (loading control). Data in A, B, D and E are representative of two or three independent biological repeats.

We detected 92 proteins that were significantly enriched in the EMCV-infected sample compared to the uninfected sample, indicating a specific interaction with MDA5 upon viral infection (Fig 1C). One of the most significantly interacting proteins was Annexin A2 (ANXA2), a multifunctional cytoplasmic protein of the Annexin family, which is involved in multiple cellular processes such as endocytosis, exocytosis, and membrane repair, but has thus far no known function in innate immune responses (*31*). ANXA2 has been reported to bind multiple different molecule types, such as phospholipids (*32*), RNA (*33*) and other proteins, with S100A10 being its most well-studied binding partner (*34*). Interaction between MDA5 and ANXA2 was confirmed by IP and direct probing for endogenous ANXA2, where the co-precipitation of ANXA2 was MDA5- and virus infection-dependent (Fig 1D). The amount of ANXA2 co-precipitated with MDA5 was small compared to the total cellular ANXA2, suggesting that only a fraction of the cellular ANXA2 either interacts with MDA5 or is able to be captured by IP. To test if the MDA5-ANXA2 interaction was mediated by RNA binding, cell lysate was treated with RNaseA, which degrades both single-stranded and double-stranded RNA, prior to IP. The ANXA2-MDA5 interaction was maintained with RNaseA treatment, suggesting a protein-protein interaction (Fig 1E).

### Direct interaction of ANXA2 and MDA5 requires the S100A10-binding domain and CARDs

To further investigate the interaction between MDA5 and ANXA2, we generated tagged constructs of ANXA2 and MDA5, either full-length or lacking specific domains (Fig 2A). When overexpressed in HEK293T cells, ANXA2 and MDA5 interacted in the absence of virus infection (Fig 2B). ANXA2 contains an S100A10-binding domain at the N-terminus, followed by four annexin repeat domains. The annexin repeats form the “core region” of the protein that mediates Ca^2+^ binding and interactions with membranes (*31*). ANXA2 constructs with sequential removal of each annexin domain (HA-ANXA2-Y269X, -Y188X, -Y109X) were successfully expressed in cells, albeit at lower expression levels compared to wild-type. Removal of all annexin repeats (HA-ANXA2-Y30X) did not yield detectable protein. Co-expression of MDA5 with HA-ANXA2-Y269X, -Y188X, -Y109X still permitted ANXA2-MDA5 interaction. However, when the S100A10 binding domain of ANXA2 was removed (HA-ANXA2-25-336), interaction with MDA5 was significantly reduced, despite equivalent protein expression levels, suggesting this domain is critical for the interaction to occur (Fig 2C).

**Figure 2.**
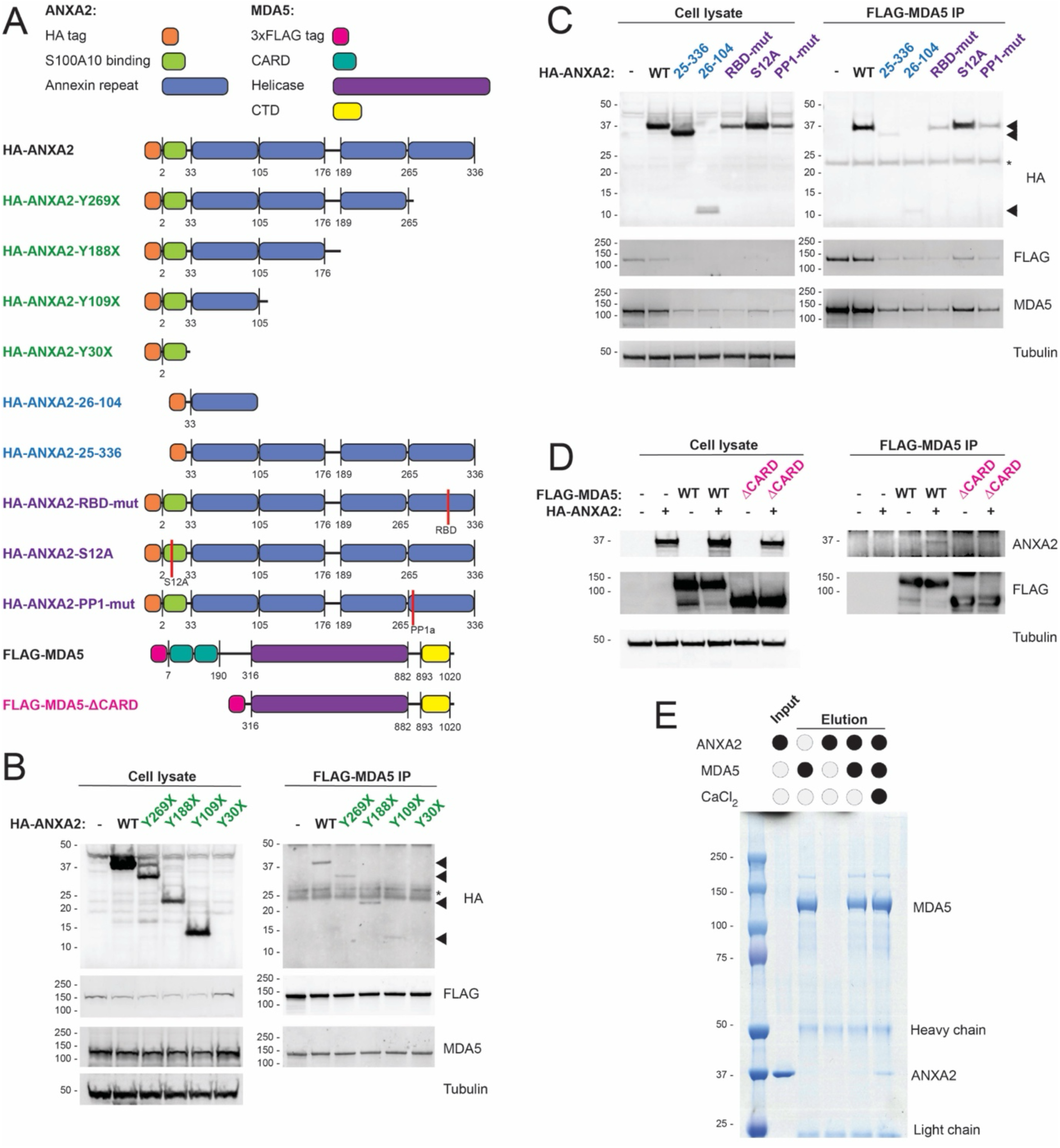
ANXA2 binds directly to MDA5 and requires intact MDA5-CARDs and the ANXA2 S100A10-binding domain. (A) Diagram of HA-ANXA2 and FLAG-MDA5 constructs containing truncations or specific site mutations. (B-D) HEK293T cells were transfected with the designated FLAG-MDA5 and HA-ANXA2 constructs and incubated for 48 hours. Cells were lysed and subjected to FLAG IP. Cell lysate (10% of IP input) and IP samples were analysed by SDS-PAGE and western blot for HA, ANXA2, FLAG, MDA5 and tubulin. Arrowheads denote HA-ANXA2 protein bands; * = IgG band. (E) Recombinant MDA5 was bound to anti-MDA5-coupled Dynabeads and incubated with recombinant ANXA2 for 30 min, in the presence or absence of 1 mM CaCl_2_. Samples were washed, eluted, resolved on SDS-PAGE and visualised by Coomassie blue staining. Lane 1 (input) is 10% of the total input. Data from (B-E) are representative of three independent biological repeats.

To explore other sites that may be important for MDA5-ANXA2 binding, we generated constructs containing mutations in the ANXA2 RNA binding domain (*33*) (HA-ANXA2-RBD-mut) and in the serine-12 phosphorylation site, which is important in the interaction with S100A10 (*35*) (HA-ANXA2-S12A). Additionally, we performed *in silico* analysis of other putative protein-interacting domains of ANXA2 using the Eukaryotic Linear Motif Prediction tool (*36*), and discovered a potential protein phosphatase-1 (PP1) domain (Fig S1). This was of interest as it has been reported that MDA5 requires dephosphorylation by PP1α for full activation (*15*). Therefore, we also generated a construct with the putative PP1 binding site mutated (HA-ANXA2-PP1-mut). The S12A mutation in ANXA2 had no effect on interactions with MDA5. The HA-ANXA2-RBD-mut and HA-ANXA-PP1-mut showed diminished binding to MDA5, though this reflects a reduced expression of these variants in cells compared to WT HA-ANXA2 (Fig 2C). Nevertheless, none of the mutations completely abolished binding to MDA5. In contrast, removal of MDA5 CARDs (FLAG-MDA5-ΔCARD), which are central mediators of the interaction with downstream adaptor MAVS, and are a site for post-translational modifications (*2*), entirely ablated interaction with ANXA2 (Fig 2D).

To ascertain whether the MDA5-ANXA2 interaction was direct or mediated via other binding partners within the cell, we generated recombinant full-length MDA5 and ANXA2. Recombinant MDA5 was immobilised on anti-MDA5 antibody-loaded beads and incubated with recombinant ANXA2 *in vitro*, followed by immunoprecipitation of MDA5. Since calcium-binding is important to ANXA2 functionality, we also tested the requirement for Ca^2+^. Direct interaction between MDA5 and ANXA2 was recapitulated *in vitro*, but only in the presence of Ca^2+^, demonstrating calcium-dependent direct binding between the MDA5 and ANXA2 in a cell-free system (Fig 2E). Thus, we demonstrated that MDA5 and ANXA2 can directly interact and that this requires the CARDs of MDA5, the S100A10-binding domain of ANXA2, and the presence of Ca^2+^.

### ANXA2 is required for MDA5 oligomerisation

To investigate the functional consequences of ANXA2 in innate immune responses, ANXA2-KO cells were generated by CRISPR/Cas9 gene editing (Fig S2). In order to assess activation of MDA5 in cells directly, we transfected an MDA5-specific RNA ligand. For this, we generated RNA extracted from EMCV-infected HeLa cells and treated with alkaline phosphatase to remove potential 5’ triphosphate structures that could activate RIG-I (EMCV-RNA). When A549 cells were transfected with EMCV-RNA, there was a robust induction of *IFNB*, *MX1* and *IFI44* in WT cells, but this was significantly reduced in ANXA2-KO cells (Fig 3A).

**Figure 3:**
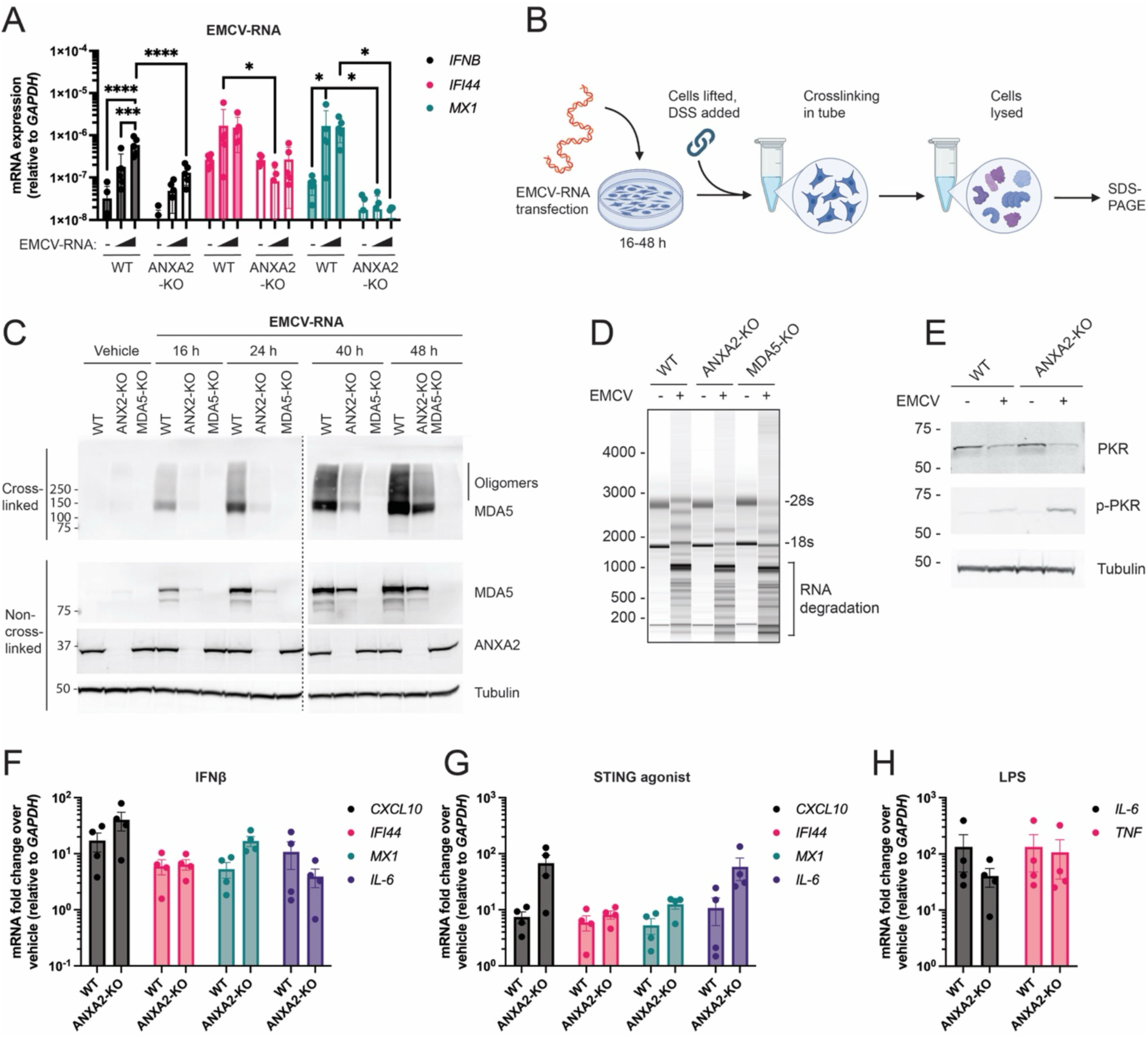
Loss of ANXA2 limits oligomerisation of MDA5 in response to EMCV-RNA, but does not affect response to other PRRs. (A) Wild-type or ANXA2-KO A549 cells were transfected with 100 ng or 400 ng per well of EMCV-RNA, or vehicle control, and incubated for 16 h. RNA was extracted and RT-qPCR performed for the indicated mRNAs. Data represent mean values of mRNA expression relative to GAPDH. (B) Schematic of MDA5 oligomerisation assay. (C) WT, ANXA2-KO or MDA5-KO A549 cells were transfected with EMCV-RNA for indicated times, prior to DSS crosslinking and lysis. Crosslinked samples were resolved on SDS-PAGE without denaturing. Non-crosslinked cell lysates were denatured with 2-mercaptoethanol prior to SDS-PAGE. Western blot for MDA5, ANXA2 and tubulin were performed. Wild-type or ANXA2-KO A549 cells were infected with EMCV (MOI = 0.01) for 24 h and (D) RNA extracted and analysed for RNA integrity, or (E) cells lysed and analysed by SDS-PAGE and western blot for PKR, phosphor-PKR (p-PKR) or tubulin. Wild-type and ANXA2-KO THP1 cells were treated for 16 h with (F) 500 U/ml of IFNΔ, (G) 20 nM STING agonist GSK3745417, (H) 1μg/ml LPS, or respective vehicle controls. RNA was extracted and RT-qPCR performed for the indicated mRNAs. Data represent mean values of fold change over vehicle control, relative to GAPDH. Data in C, D and E are representative images of two or three independent biological repeats. Data points in A and F-H are from four or five independent biological repeats. Statistical significance was tested by 2-way ANOVA with Sidak’s multiple comparison test.

A key step in the activation of MDA5 is the oligomerisation of multiple MDA5 monomers on dsRNA, which promotes clustering of CARDs and thus downstream signalling via MAVS (*14*). We developed an MDA5 oligomerisation assay using disuccinimidyl suberate (DSS) crosslinking of intact cells, followed by lysis and non-reducing SDS-PAGE to visualise MDA5 oligomers by Western blot (Fig 3B). Transfection of A549 cells with EMCV-RNA promoted robust time-dependent MDA5 oligomerisation, visible as a high molecular weight smear (Fig 3C). MDA5 oligomerisation was detected at 16 h post-transfection in WT cells and increased over time. Comparatively, ANXA2-KO cells showed absent or delayed MDA5 oligomerisation. These results demonstrate that ANXA2 promotes MDA5 activity via enhancing its oligomerisation, a crucial step in the signalling pathway.

### ANXA2 does not affect activation of other PRRs

Long dsRNA is the established agonist for MDA5, but dsRNA can also activate other innate immune sensors such as PKR and the OAS-RNase L pathway, which induce global protein translation shutdown and RNA degradation, respectively (*37*). We wanted to determine if ANXA2 influenced these dsRNA-binding receptors. Activated PKR is phosphorylated, whereas RNase L activation can be demonstrated by decrease of 28S and 18S ribosomal RNAs and increase in cellular RNA degradation products. EMCV infection induced both cellular RNA degradation and PKR phosphorylation (and concomitant reduction in unphosphorylated PKR), and neither was impeded by loss of ANXA2 (Fig 3D,E).

To determine if loss of ANXA2 could affect other innate immune signalling pathways in cells, we tested the response of WT and ANXA2-KO THP1 cells to additional stimuli. The response to IFNΔ treatment was equivalent between the two genotypes (Fig 3F), indicating there is no defect in the transcriptional response to interferon in ANXA2-KO cells. The treatment of cells with the STING agonist GSK3745417, which induces both a T1-IFN response and an NF-κB-mediated proinflammatory response, was also equivalent between the two genotypes, although there was a trend towards ANXA2-KO cells showing a potentiated response to STING activation (Fig 3G). Similarly, there was no difference in the response to LPS, a TLR4 agonist known to strongly induce NF-κB-mediated IL-6 and TNF expression (Fig 3H). Overall, loss of ANXA2 did not prevent the induction of T1-IFN or NF-κB-mediated gene signatures in response to other PRR ligands, indicating there was no defect in activation of these transcriptional pathways in the absence of ANXA2.

### ANXA2 is required for type I interferon response to RNA virus infection

To investigate the role of ANXA2 in response to virus infection, we infected WT, ANXA2-KO or MDA5-KO THP1 cells with EMCV and assessed transcriptional changes. Loss of ANXA2 in THP1 cells inhibited the T1-IFN response to EMCV infection, seen by a significant reduction in the expression of *IFNB* and the ISGs *CXCL10, IFI44, IFIT1* and *MX1* (Fig 4A). This reduced response occurred despite equivalent levels of viral RNA (EMCV 5’NTR) in cells, indicating the virus replicated equally in ANXA2-KO cells compared to wild-type (Fig 4B). Transcriptomic analysis of THP1 cells infected with EMCV showed that ANXA2-KO cells had significantly fewer gene expression changes compared to wild-type (Fig 4C, Fig S3A). The THP1 response to EMCV was MDA5-dependent, since loss of MDA5 abolished the transcriptional response to EMCV. Comparison of the gene expression changes in ANXA2-KO vs. WT in response to EMCV showed that nearly all gene changes in ANXA2-KO were also affected in WT, whereas there were 470 genes exclusively upregulated in WT but not in ANXA2-KO (Fig 4C, bottom right panel), indicating the response in ANXA2 was of a significantly lower magnitude. Interferon response and inflammation gene sets were upregulated in response to EMCV in both WT and ANXA2-KO cells, but this was lower in the ANXA2-KO and significantly diminished in MDA5-KO cells (Fig 4E and Fig S4A,B). It is notable that ANXA2-KO THP1 cells showed a mild but detectable increased basal T1-IFN signature in uninfected cells, compared to WT uninfected cells (Fig 4D,E). Nevertheless, despite this increased basal inflammatory profile, ANXA2-KO cells did not respond as strongly to infection as wild-type cells.

**Figure 4:**
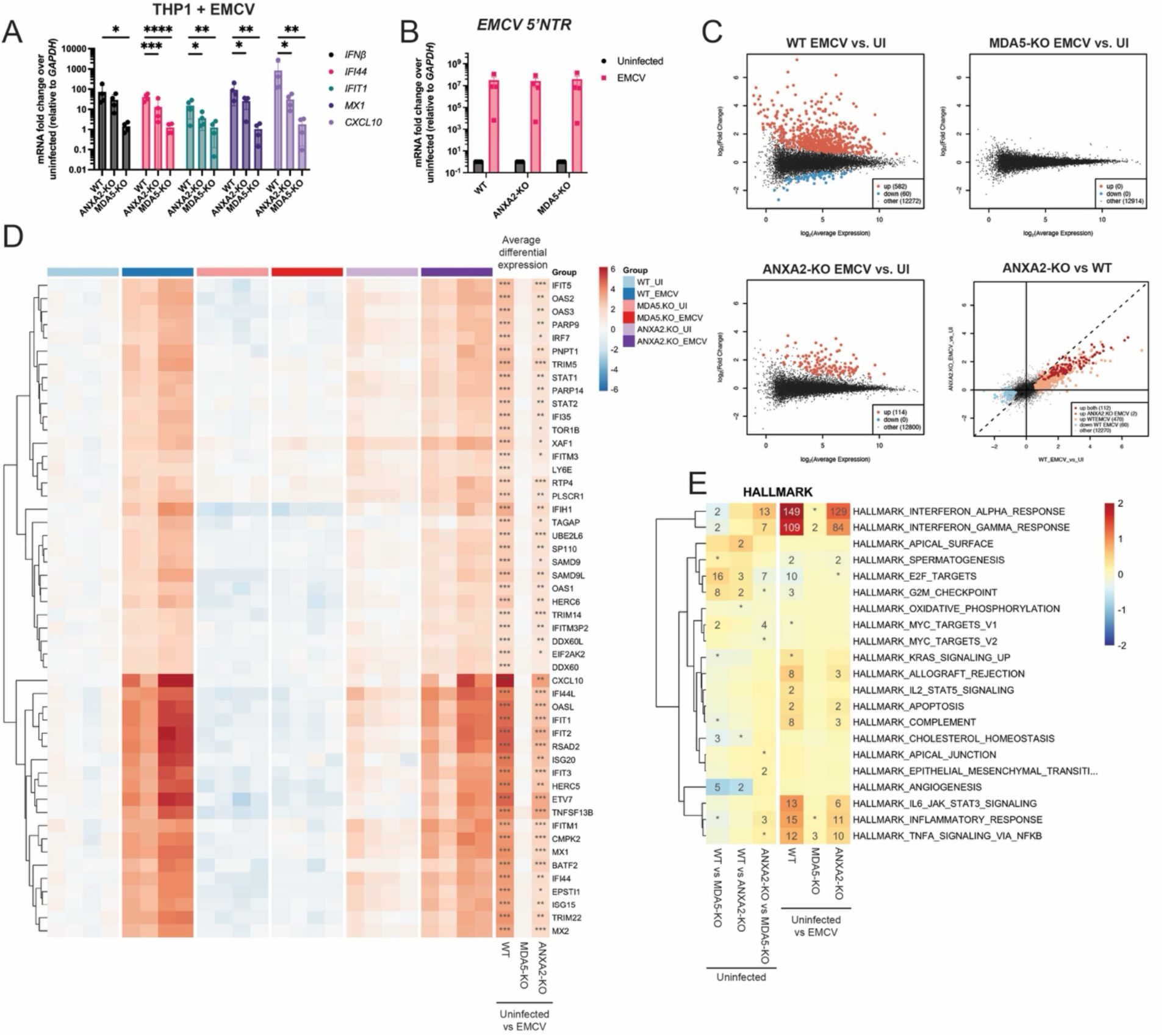
Loss of ANXA2 dampens THP1 cell response to EMCV. Wild-type, ANXA2-KO and MDA5-KO THP1 cells were infected with EMCV (MOI = 5) for 22 hours. RNA was extracted and RT-qPCR performed for (A) indicated mRNAs and (B) EMCV 5’ non-translated region (NTR), or (C-F) analysed by RNA-seq. Data in (A-B) are shown relative to *GAPDH*, and data in (A) represents fold change over uninfected. Bars in (A-B) indicating the mean values from four independent biological repeats. Statistical significance was tested by 2-way ANOVA with Sidak’s multiple comparison test. (C) Differential expression is depicted as log_2_ fold changes of all genes between infected (EMCV) and uninfected (UI) samples for each genotype versus average gene expression or between ANXA2-KO and WT infected cells directly. (D) Heat map of relative gene expression of all samples compared to WT uninfected for genes changed in response to infection (scale truncated to ±6). Average log_2_ fold changes between infected and uninfected samples are indicated in right three columns. Asterisks indicate genes significantly different between comparisons (* p < 0.05, ** p < 0.01 and *** p < 0.001). (E) Gene set test results for Hallmark gene set collections. Colours indicate the average log_2_ fold changes of all genes in the gene set (scale truncated to ±2). From left to right: columns 1-3 compare between genotypes in uninfected cells, columns 4-6 compare between infected and uninfected cells within each genotype. The significance of each gene set is indicated by the text as the −log_10_ FDR-adjusted p-value threshold (* denotes adjusted p < 0.05, 2 denotes p < 0.01, etc.).

To extend our investigations into the role of ANXA2 in antiviral innate immune signalling, we analysed the response to West Nile virus (WNV; Kunjin subtype), which is known to activate both MDA5 and RIG-I (*38–40*). THP1 cells were infected with WNV for 24 h and analysed for select T1-IFN response genes and total transcriptomic changes. In contrast to EMCV infection, where loss of ANXA2 blunted T1-IFN responses, the T1-IFN response to WNV infection was almost entirely ANXA2-dependent in THP1 cells (Fig 5A, Fig S3B). There was a small but significant reduction in levels of WNV non-structural protein 1 (NS1) RNA in ANXA2-KO cells, and an increase in MDA5-KO cells, compared to wild-type (Fig 5B). Strikingly, the transcriptional response to WNV was largely ablated in the ANXA2-KO cells (Fig 5C). The effect of ANXA2 loss in the response to WNV was more drastic than that of MDA5 loss (Fig 5C,D), supporting the concept that T1-IFN response to WNV is not only dependent on MDA5, but also on other innate immune sensors. Transcriptional response to WNV, similarly to responses to EMCV, consisted of the induction of interferon and inflammation gene sets (Fig 5E and Fig S4C,D), which were significantly reduced in ANXA2-KO (Fig 5D, right column). We also performed WNV infections in WT, ANXA2-KO and MDA5-KO A549 cells, and found that, similar to THP1 cells, the innate responses were largely dependent on ANXA2 (Fig S3C, Fig S4C, Fig S5).

**Figure 5:**
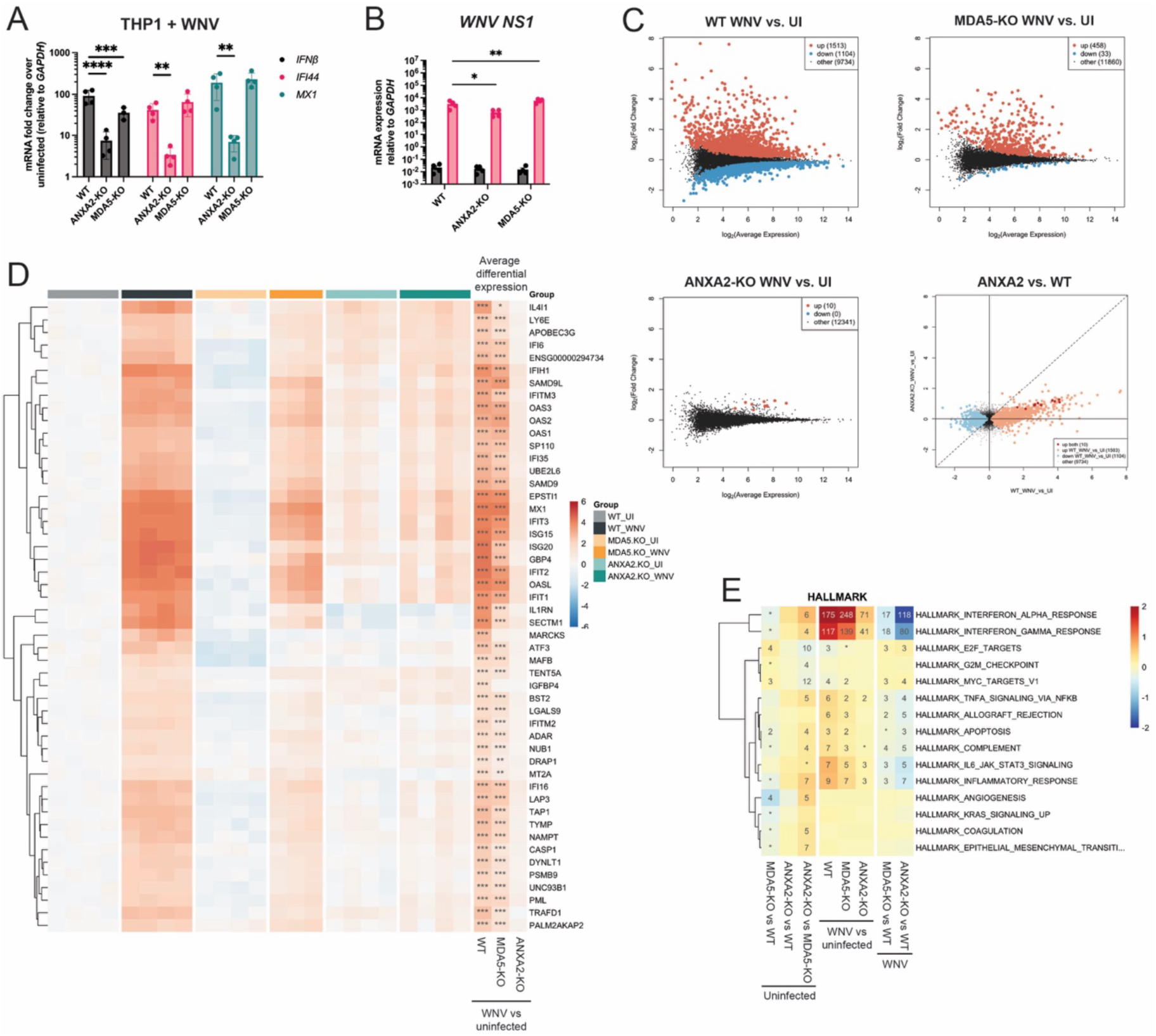
Loss of ANXA2 ablates THP1 cell response to WNV. Wild-type, ANXA2-KO and MDA5-KO THP1 cells were infected with WNV (MOI = 1) for 24 hours. RNA was extracted and RT-qPCR performed for (A) indicated mRNAs and (B) WNV NS1 RNA, or (C-F) analysed by BRB-seq. Data in (A-B) are shown relative to GAPDH, and data in (A) represents fold change over uninfected. Bars in (A-B) indicating the mean values from four independent biological repeats. Statistical significance was tested by 2-way ANOVA with Sidak’s multiple comparison test. (C) Differential expression is depicted as log_2_ fold changes of all genes between infected (WNV) and uninfected (UI) samples for each genotype versus average gene expression or between WT and ANXA2-KO cells in infected cells directly. (D) Heat map of relative gene expression of all samples compared to WT uninfected for genes changed in response to infection. Average log_2_ fold changes between infected and uninfected samples are indicated in right three columns. Asterisks indicate genes significantly different between comparisons (* p < 0.05, ** p < 0.01 and *** p < 0.001). (E) Gene set test results for Hallmark gene set collections. From left to right: columns 1-3 compare between genotypes in uninfected cells, columns 4-6 compare between infected and uninfected cells within each genotype, and columns 7-8 compare between genotypes in infected cells. Colours indicate the average log_2_ fold changes of all genes in the gene set (scale truncated to ±2). The significance of each gene set is indicated by the text as the −log_10_ FDR-adjusted p-value threshold (* denotes adjusted p < 0.05, 2 denotes p < 0.01, etc.).

### ANXA2 promotes the innate response to WNV in human neuronal cells

WNV infection can, in some cases, spread to the brain and lead to life-threatening encephalitis. WNV is known to replicate in neuronal cells and induce cytopathic effects (*41*). To assess the role of ANXA2 in response to virus infection in a more disease-relevant cell model, we generated human neuronal progenitor cells (NPC) from induced pluripotent stem cells (iPSC). NPCs were successfully differentiated from WT and ANXA2-KO iPSCs, as indicated by upregulation of specific markers *NESTIN*, *PAX6* and *TUBB3* (Fig6A). NPC were readily infected by WNV, indicated by the presence of viral NS1 RNA (Fig 6B). WNV infection induced a robust T1-IFN response, as shown by upregulation of *IFNA* and ISGs (Fig 6B). However, this response was lower in ANXA2-KO cells compared to WT, indicating that ANXA2 is required for robust response to WNV in human neuronal cells.

**Figure 6:**
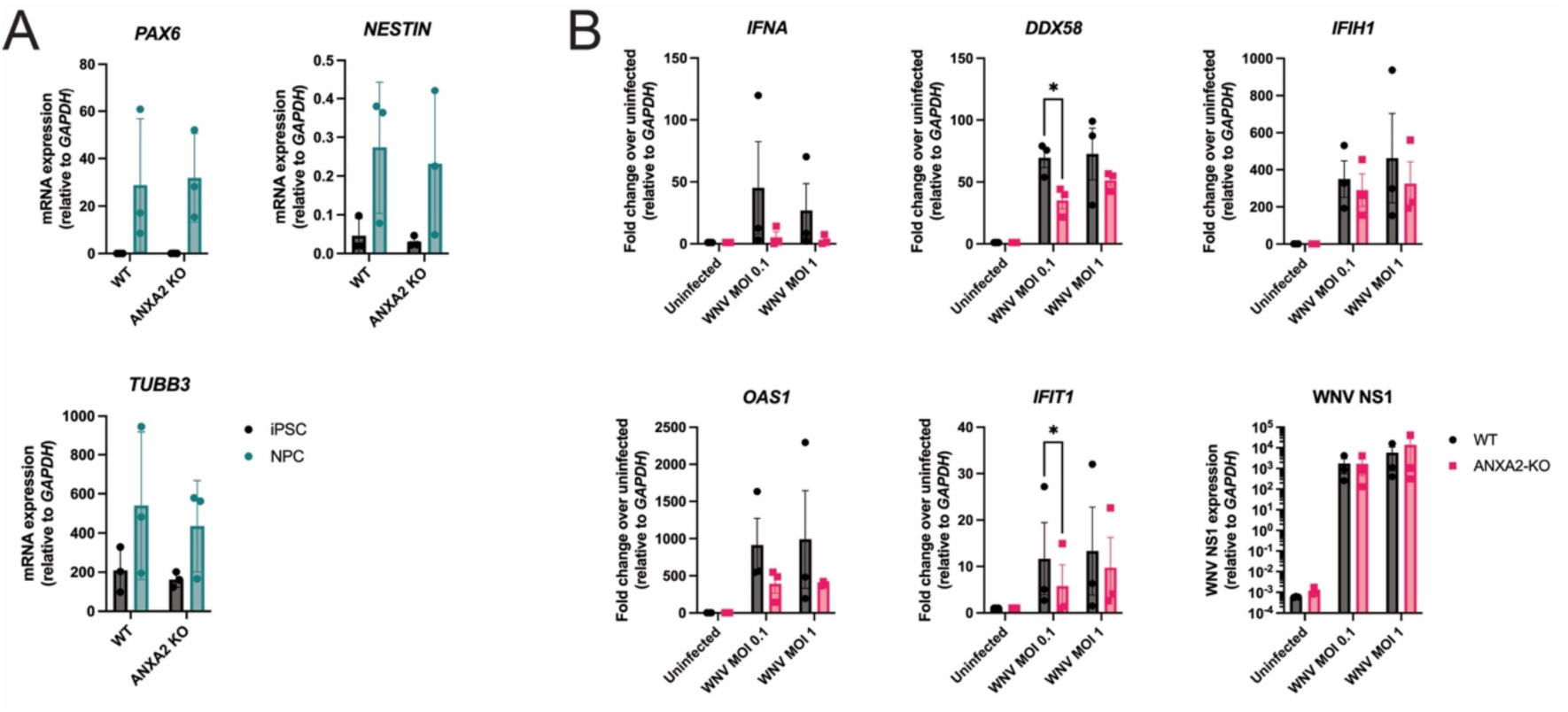
Loss of ANXA2 affects response to WNV in human neuronal progenitor cells. (A) RNA was extracted from WT or ANXA2-KO human iPSC and iPSC-derived NPCs and RT-qPCR performed for the indicated NPC-associated mRNAs. Data represents fold-change over iPSC, relative to *GAPDH*. (B) WT and ANXA2-KO NPC were infected with WNV at the indicated MOI for 24 or 48 h. RNA was extracted and RT-qPCR performed for the indicated mRNAs. Data shown is mRNA expression relative to *GAPDH*. Data in A and B are mean values from three independent biological repeats. Statistical significance was tested by 2-way ANOVA with Sidak’s multiple comparison test. Data in B are representative images from two independent biological repeats.

### ANXA2 mediates RIG-I activation

Given the drastic effect of loss of ANXA2 in cellular responses to WNV infection, and since WNV can activate both RIG-I and MDA5 (*40*), we hypothesized that ANXA2 may also mediate RIG-I activation. To test this, we assessed WT and ANXA2-KO A549 cell response to the well-defined RIG-I ligand 3p-hpRNA. We found significantly reduced T1-IFN response to transfected 3p-hpRNA in ANXA2-KO cells compared to WT (Fig 7A), though some ISG levels were unchanged. Influenza A virus (IAV) is a potent inducer of RIG-I, but not known to stimulate MDA5 (*30, 42*). To assess the role of ANXA2 in response to IAV, we first extracted RNA from IAV-infected MDCK cells (IAV-RNA) and transfected the RNA into A549 cells. This approach bypasses host restriction by virus-encoded proteins that can inhibit innate immune sensing (*42*). We found that IAV-RNA strongly stimulated WT cells but this was significantly reduced in ANXA2-KO (Fig 7B). To examine the response to live IAV infection in A549 cells, we used a strain of IAV lacking the viral NS1 protein (IAV-ΔNS1), as NS1 is a suppressor of T1-IFN responses (*43*). Thus, use of the IAV-ΔNS1 enables detection of the RIG-I pathway activation in response to infection. A549 cells were successfully infected with IAV-ΔNS1, as indicated by expression of IAV M1 viral RNA (Fig 7C), and showed a T1-IFN response that was partially reduced in the absence of ANXA2. Combined, these data suggest that ANXA2 may also be involved in promoting the RIG-I pathway (Fig 7D).

**Figure 7:**
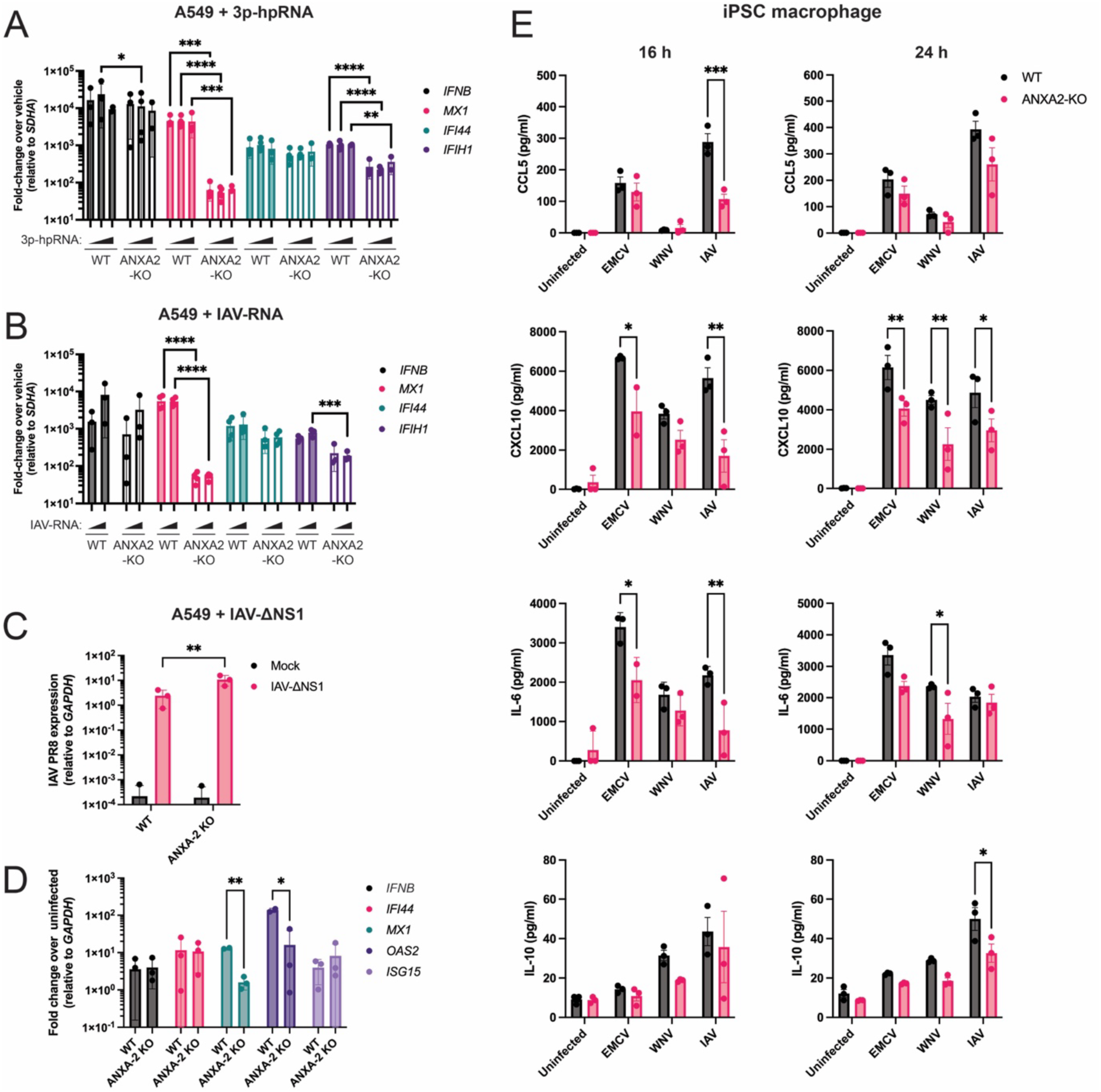
ANXA2 is required for robust RIG-I responses. A549 cells were transfected with (A) 12.5, 25 or 50 ng of 3p-hpRNA, (B) 100 or 400 ng of IAV-RNA, or vehicle control (lipofectamine only) for 16 h. RNA was extracted and RT-qPCR performed for the indicated mRNAs. Wild-type or ANXA2-KO A549 cells were infected with NS1-deleted influenza virus (IAV ΔNS1; MOI = 10) for 16 h. RNA was extracted and RT-qPCR performed for (C) IAV M1 RNA and (D) the indicated mRNAs. (E) WT or ANXA2-KO human iPSC-derived macrophages were infected with EMCV, WNV or IAV (MOI = 10) for 16 or 24 hr and released cytokines were measured. Data in (A, B and D) represent mean fold-change over vehicle or uninfected control, relative to *SDHA* from three or four independent biological repeats. Data in C represents expression relative to *GAPDH*. Data in (E) represent mean of three independent biological repeats. Statistical significance was tested by 2-way ANOVA with Sidak’s multiple comparison test.

Macrophages are innate immune cells that play a critical role in the early response to infection and orchestrate the immune response in multiple ways (*44*). In the lung, alveolar macrophages are a major source of pro-inflammatory cytokines produced during IAV infection (*45*). Monocytes/macrophages are also target cells in WNV infection (*39, 46*). Thus, to study the role of ANXA2 in this crucial immune cell type, we generated human macrophages from WT and ANXA2-KO iPSCs. Successful differentiation into macrophages was evident by high expression of macrophage markers CD14, CD163, CD16 and CD11b and no expression of hematopoietic stem cell marker CD34 (Fig S6). We infected the iPSC-derived macrophages with EMCV, WNV and IAV and assessed the inflammatory cytokines released into the cell culture supernatant. We found consistently that there was a lower response to virus infection in the ANXA2-KO cells compared to WT (Fig 7E). This was significant for IAV infection, in particular at the earlier 16 h time point, with lower levels of the proinflammatory cytokines CCL5, CXCL10 and IL-6. Similarly, release of CXCL10 and IL-6 in response to EMCV was also significantly reduced. The response to WNV at 16 h was low, likely due to the slower replication cycle of this virus. However, at 24 h there was significantly less CXCL10 and IL-6 by in the ANXA2-KO cells infected with WNV, compared to WT cells. Taken together, we find that ANXA2 is necessary for a robust innate immune response to several viruses, and in multiple disease-relevant cell types, by promoting RLR signalling in response to infection.

## Discussion

We report here that ANXA2 plays a critical role in the RLR pathways. Using multiple cell types and virus infections, we found that loss of ANXA2 significantly dampens RLR pathway activation, but has no effect on other PRR pathways that utilise similar downstream mediators, e.g. STING and TLR4. This demonstrates that the role of ANXA2 is upstream of transcription factor activation and gene expression. Furthermore, ANXA2 does not play a role in other dsRNA-binding PRRs, such as PKR and OASs, which indicates it has a function specific to RLRs.

We have multiple lines of evidence for direct protein-protein interactions between MDA5 and ANXA2. When overexpressed in cells, ANXA2 co-precipitated with MDA5, and this interaction was dependent on the CARDs of MDA5 and the S100A10-binding domain of ANXA2. At endogenous levels, however, the MDA5-ANXA2 interaction only occurred in response to viral infection, suggesting it is a specific ligand-triggered event. *In vitro* experiments using full-length recombinant proteins recapitulated binding between MDA5 and ANXA2, indicating these proteins do not require additional protein factors for their interaction. However, *in vitro* interactions were dependent on Ca^2+^. ANXA2 has multiple Ca^2+^ binding sites, and binding of Ca^2+^ regulates ANXA2 interactions with other molecules, proteins, and RNA (*47, 48*). There have been reports that virus infections, such as poliovirus (*49*), SARS-CoV-2 (*50*), rotavirus (*51, 52*), and WNV (*53*) can induce an increase in intracellular Ca^2+^ levels. Changes to intracellular Ca^2+^ levels could thus represent an additional mechanism of regulation of RLR signalling, where an increase in Ca^2+^ associated with infection may promote ANXA2 interaction with RNA and/or MDA5, to enhance receptor activation. Future investigations are needed to determine the role of Ca^2+^ levels in regulation of ANXA2-MDA5.

In addition to MDA5, data from infection with WNV and IAV indicate that ANXA2 is also required for robust RIG-I activation. However, experiments using transfected RIG-I RNA ligands showed that RIG-I activation was not entirely dependent on ANXA2. This discrepancy between the requirement for ANXA2 in cellular responses to active viral infection vs stimulation with a purified RIG-I ligand might be indicative of ANXA2 playing a specialised role in RLR pathways that is bypassed when the ligand is directly provided to the cytosol at high concentrations. We can speculate that ANXA2 may support the co-localization of RLRs within relevant subcellular compartments that promote interactions between receptors and RNA ligands, as has been demonstrated for ZCCHC3 (*28, 54*). ZCCHC3 was shown to promote RLR and cGAS signalling via phase separation with nucleic acids, thus enriching the nucleic acid ligand within a distinct subcellular location to increase interaction with PRRs. The pleiotropic functions of ANXA2, particularly in the movement of proteins and organelles within the cell, would support this hypothesis. Another Annexin family member, ANXA11, was recently shown to promote phase separation in neurons, enabling interactions between RNA granules and lysosomes (*55*). Indeed ANXA2, like other Annexins, is recruited to lysosomes to promote their repair (*56*). Whether ANXA2 similarly supports phase separation or subcellular distribution of dsRNA and/or RLRs to promote RLR activation is yet to be investigated.

ANXA2 has been shown to play a role in numerous cellular functions, including endocytosis, exocytosis, lysosomal repair, and membrane organisation (*34, 47*). Additionally, several viruses employ ANXA2 as an entry receptor for cellular invasion (*57*). However, the viruses studied here did not require ANXA2 for cellular entry, as our ANXA2-KO cells were equally permissible to viral replication compared to WT cells. Thus, we revealed a novel function of ANXA2 in directly interacting with MDA5 and promoting receptor oligomerisation. This is similar to the recent report that ADAM9, a metalloprotease, promotes MDA5 oligomerisation and plays an essential role in protection from EMCV *in vivo* (*27*). The high expression level of ADAM9 in fibroblasts likely supports this protective role for ADAM9 in the heart, a target organ of pathology for EMCV. The chaperone-like protein 14-3-3η was also reported to promote MDA5 oligomerisation in cells in response to Poly(I:C) and EMCV infection (*24*) and is directly antagonised by Zika virus in order to evade RLR sensing (*25*). ANXA2 is highly expressed in monocytes and epithelial cells, and functional redundancy exists between different Annexins. It is possible that we see a critical role for ANXA2 in the RLR pathway in the cells studied here because ANXA2 is highly expressed in these cell types. Whether other Annexins play similar roles in different cell types, depending on their abundance, is yet to be determined. It is possible that multiple chaperones/mediators of RLR activation function concurrently or in sequence to promote complete pathway activation within one cell. Alternatively, the dependence on different mediators, be they ANXA2 in monocytes or ADAM9 in fibroblasts, may be cell-type dependent.

We found that loss of ANXA2 in THP1 cells also resulted in increased basal T1-IFN signature, in the absence of infection. It is possible that ANXA2 plays additional roles in maintenance of tonic T1-IFN expression, via a yet undescribed mechanism. Indeed, it has been reported that ANXA2-knockdown in renal tubular epithelial cells caused an increased inflammatory gene signature (*58*). However, this was not evident in A549 lung epithelial cells. The status of basal MDA5 expression could be a factor, as MDA5 is lowly expressed in A549 cells in the absence of infection or T1-IFN treatment (Fig S1). Alternatively, this could be due to differences in basal expression of regulators of interferon signalling, e.g. USP18 (*59*) or SOCS proteins (*60*).

The dependency of ANXA2 for cellular detection of WNV was striking. Flaviviruses are known to replicate inside membraned replication compartments at the cellular endoplasmic reticulum. This process shields the viral replicating RNA, which contains RNA species that can activate both RIG-I and MDA5, from the cytosolic sensors. Recently, it was shown that in a small proportion of cells, negative-sense single-stranded viral RNA from WNV escapes replication compartments and activates RIG-I (*40*). However, the mechanism of escape of this viral RNA, and whether it is driven by host factors, remains undetermined. ANXA2 has a known function in processes involved in joining of membranous compartments. It would be interesting to investigate if ANXA2 may play a role in the exposure of viral RNA from replication compartments to promote sensing of viral RNA.

## Materials and Methods

### Cell culture and virus production

Cells were cultured at 37°C and 5% CO_2_ and routinely screened for mycoplasma contamination. THP1 and BHK-21 cells were maintained in RPMI, Vero, HeLa, MDCK, p125HEK (*61*), and A549 cells were maintained in DMEM, both supplemented with 10% v/v foetal calf serum (FCS), 100 U/ml Penicillin-Streptomycin and 2 mM L-glutamine (all reagents Gibco). THP1 cells were differentiated into macrophage-like cells prior to experiments by 24 h treatment with 10 ng/ml phorbol myristate acetate (PMA; Invivogen). For generation of ANXA2 knockout cells using CRISPR/Cas9 technology, two sgRNAs targeting upstream and downstream of exon 2 of the human ANXA2 gene were cloned into either pX458-eGFP or pX458-Ruby (Addgene 110164, deposited by Dr. Philip Hublitz), as described previously (*61*). LTX Transfection kit (ThermoFisher) was used to transfect 4 x 10^6^ THP1 cells with 10 μg of both plasmids, or 7 x 10^5^ A549 or p125HEK cells with 1 μg of both plasmids according to manufacturer’s instructions. After 24 hr, mRuby- and eGFP-double positive cells were single-cell sorted into 96-well plates containing 50% fresh medium and 50% filtered conditioned medium from parental cells. Surviving clones were expanded and screened by immunoblotting for loss of ANXA2 expression, and Sanger sequenced to ascertain mutation at the gene locus. EMCV was produced in BHK-21 cells, and harvested from supernatant after centrifugation at 4,000 xg and 0.4 μm filtration, before single-aliquot freezing at −80°C. Viral content was quantified by plaque assay on BHK-21 cells. WNV (Kunjin subtype, gift from J. Mackenzie, University of Melbourne) was produced in Vero cells, and harvested from supernatant by centrifugation at 4,000 xg and 0.2 μm filtration before single-aliquot freezing at −80°C. Viral content was quantified by plaque assay on Vero cells. The IAV strain HKx31 (H3N2) was grown in 10-day embryonated chicken eggs (kindly provided by P. Reading, University of Melbourne) and titrated on MDCK cells. IAV PR8 virus lacking the NS1 gene (referred to as IAV-ΔNS1; a kind gift from A. Garcia-Sastre (*43*)) was propagated as described previously (*62*).

### Antibodies and Reagents

MDA5 antibodies were generated by C. Song and B. Jin. Clone 16 (herein referred to as mAb 16) and clone 17 (herein referred to as mAb 17) were used for immunoprecipitation and immunoblotting, respectively. Antibodies were purified from hybridoma cell culture supernatants by the Monash Antibody Discovery Platform using Protein A. Antibodies against ANXA2 (Cell Signalling Technology, 8235), PKR (Cell Signalling Technology, 12297), phospho-PKR (Abcam, ab32036), tubulin (Abcam, ab6160), FLAG (M2, Monash Antibody Discovery Platform) and HA (12CA5, Monash Antibody Discovery Platform) were used for immunoblotting at 1:1000. Secondary antibodies AlexaFluor680 anti-mouse (Invitrogen, A21057), AlexaFluor680 anti-rabbit (Invitrogen, A21088), Dylight800 anti-mouse (Cell Signalling Technology, 5257P), and IRDye800 anti-rabbit (Li-Cor, 925-32211) were used at 1:6000.

### Generation of protein expression constructs for mammalian expression

All ANXA2 constructs were cloned into pcDNA3.1, contained an HA tag at the N-terminus, and were generated by GenScript. For truncated mutants HA-ANXA2-Y269X, - Y188X, -Y109X and -Y30X, the tyrosine was mutated to a stop codon to induce early termination. For the PP1 mutant, alanine substitutions were made at positions K266, L268, and F270. For the RNA binding domain mutant, serine substitutions were made at positions K307, R308, K309, K312, Y316 and Q320. To generate the FLAG-MDA5 construct, the MDA5 mRNA sequence was PCR amplified from cDNA isolated from HEK293 cells treated with IFN, and inserted into pcDNA3.1-TOPO (Invitrogen) containing 3xFLAG at the N-terminus. The FLAG-MDA5-ΔCARD construct was generated by PCR amplification of bases 295-1025 from the FLAG-MDA5 gene, and insertion into pcDNA3.1-TOPO. All constructs were verified by whole plasmid sequencing.

### MDA5 purification

The DNA sequence encoding MDA5 was derived from cDNA isolated from K562 cells and cloned into a modified 438-C vector (*63*) with an N-terminal hexahistidine-MBP tag followed by a 3C protease cleavage site. Bacmids were isolated after transformation of DH10EMBacY cells (Geneva Biotech) with isolated plasmid DNA. Bacmid DNA was transfected into Sf9 cells using FuGENE 6 reagent (Promega) to generate recombinant baculovirus, which was further amplified in Sf9 cells. For MDA5 expression, Hi5 cells were infected with baculovirus and incubated for 72 hours. Unless otherwise stated, next steps were performed at 4°C. Cells were harvested by centrifugation and lysed by sonication in MDA5 lysis buffer (50 mM sodium phosphate-NaOH, 400 mM NaCl, 20 mM imidazole, 20% [v/v] glycerol, 1 mM DTT; pH 6.0 at 25 °C) supplemented with protease inhibitors (1 mM PMSF, 2 mM benzamidine, 1 µM leupeptin and 2 µM pepstatin). MDA5 was purified by affinity chromatography using a HisTrap HP column (Cytiva) equilibrated in MDA5 lysis buffer. MDA5 was washed with a high salt wash (MDA5 lysis buffer containing 2 M NaCl and 30 mM imidazole), an ATP wash (MDA5 lysis buffer containing 100 mM NaCl, 3 mM MgCl_2_, 3 mM ATP and 30 mM imidazole), and eluted using an imidazole gradient ranging from 20 mM to 500 mM. MDA5 was cleaved by 3C protease and dialysed overnight against buffer C (50 mM sodium phosphate-NaOH, 200 mM NaCl, 10% [v/v] glycerol; pH 6.0 at 25 °C). The following day, MDA5 was diluted to a final NaCl concentration of 100 mM and purified using a HiTap Heparin HP column (Cytiva) equilibrated in buffer A (50 mM sodium phosphate-NaOH, 100 mM NaCl, 10% [v/v] glycerol, 1 mM DTT; pH 6.0 at 25 °C) and eluted using a gradient ranging from 100 mM to 2 M NaCl. MDA5-containing fractions were pooled and further supplemented with 40 mM imidazole, then purified by HisTrap HP. Flow through was concentrated using 30 kDa concentrators (Amicon) and purified using a Superdex 200 10/300 column (Cytiva) equilibrated in MDA5 storage buffer (50 mM BisTris-HCl, 150 mM NaCl, 10 % [v/v] glycerol; pH 6.5 at 25 °C). Protein concentration and the ratio of absorbance at 260 nm and 280 nm (to confirm a lack of RNA contamination) were measured using UV absorbance in a quartz cuvette with an optical path of 1 cm.

### ANXA2 purification

ANXA2 was amplified by PCR from the above-described pcDNA vector with ANXA2 insertion and cloned into a pOPINB vector with an N-terminal hexahistidine tag and 3C protease cleavage site. ANXA2 was expressed in *E. coli* BL21(DE3)RIL in LB supplemented with 50 mg/ml kanamycin at 37 °C. Expression was induced at OD600 between 0.6-0.8 by addition of 1 mM IPTG for 18 hours at 18 °C. Unless otherwise stated, the following steps were performed at 4°C. Cells were harvested by centrifugation, resuspended in ANXA2 lysis buffer (50 mM HEPES-NaOH, 200 mM NaCl, 20 mM imidazole, 20 % [v/v] glycerol, 1 mM DTT; pH 8 at 25 °C) supplemented with 0.1% (v/v) IGEPAL and protease inhibitors (1 mM PMSF, 2 mM benzamidine, 1 µM leupeptin and 2 µM pepstatin), and lysed by sonication. ANXA2 was purified by affinity chromatography using a HisTrap HP column (Cytiva) equilibrated in ANXA2 lysis buffer. After loading, the column was washed with ANXA2 lysis buffer containing 1 M NaCl and 30 mM imidazole and eluted using an imidazole gradient ranging from 20 mM to 750 mM. ANXA2 was cleaved by 3C protease and dialysed overnight against buffer D (50 mM HEPES-NaOH, 50 mM NaCl, 10 % [v/v] glycerol; pH 8 at 25 °C). The next day, ANXA2 was supplemented with 5 mM CaCl_2_ and purified using a HiTrap Heparin HP column (Cytiva) equilibrated in buffer E (50 mM HEPES-NaOH, 50 mM NaCl, 5 mM CaCl_2_, 10 % [v/v] glycerol, 1 mM DTT; pH 8 at 25 °C). Protein was eluted using a gradient ranging from 50 mM to 1 M NaCl. ANXA2-containing fractions were further supplemented with 20 mM imidazole and purified by HisTrap HP. Flow through was concentrated using 10 kDa concentrators (Amicon) and purified using a Superdex 75 10/300 column (Cytiva) equilibrated in ANXA2 storage buffer (50 mM HEPES-NaOH, 150 mM NaCl, 10 % [v/v] glycerol; pH 8 at 25 °C). Protein concentration and the ratio of absorbance at 260 nm and 280 nm (to confirm a lack of RNA contamination) was measured using UV absorbance in a quartz cuvette with an optical path of 1 cm.

### Immunoblotting

Cells were lysed with lysis buffer (10 mM Tris, 50 mM NaCl, 30 mM sodium pyrophosphate, 50 mM NaF, 5 µM ZnCl_2_, 0.5% IGEPAL and complete protease inhibitor cocktail (Roche)) and protein was quantified by BCA assay (Pierce). Samples were denatured using NuPAGE 4x LDS sample loading buffer (Life Technologies) and 10% 2-mercaptoethanol, and heating at 95°C for 5 min. Samples were resolved by electrophoresis on 4-12% Bis-Tris gels with MOPS Running Buffer (Life Technologies NuPAGE system) and transferred to nitrocellulose membrane by electrophoresis at 100V for 2 h in cold transfer buffer (Life Technologies) with 10% methanol. Membranes were blocked with Intercept Blocking Buffer (Li-Cor) for 1 hr, and probed with primary (overnight at 4°C) and secondary (1 h at room temperature) antibodies diluted in Intercept Blocking Buffer, with rotation. Membranes were washed four times in 0.05% IGEPAL in Tris-buffered saline (50 mM NaCl, 50 mM Tris-HCl, pH 7.6) for 5 min each wash after each antibody incubation. Membranes were analysed on Bio-Rad ChemiDoc.

### Immunoprecipitation

Cell lysate was prepared as above and quantified by BCA assay. For immunoprecipitation of endogenous MDA5, anti-MDA5 mAb 16 was covalently coupled to Dynabeads (Invitrogen, 14311D) according to manufacturer’s instructions, using 2 mg Dynabeads and 40 µg of mAb 16 per sample. mAb 16 antibody-coupled Dynabeads were washed with lysis buffer, and incubated with 1 mg of cell lysate for 3 h at 4°C with rotation. Samples were washed three times with Wash Buffer (10 mM Tris, 50 mM NaCl, 30 mM sodium pyrophosphate, 50 mM NaF, 5 µM ZnCl_2_) containing 0.5 % NP-40 (IGPAL), and once with Wash Buffer without NP-40. Samples were eluted using NuPAGE 4x LDS sample buffer (Invitrogen) and heating at 70°C for 10 min. Samples were denatured and resolved by SDS-PAGE and western blotting as above.

For immunoprecipitation of FLAG-MDA5, 500 µg of cell lysate was pre-cleared by incubating with 20 µl of Protein G Dynabeads (ThermoFisher) for 1 h at 4°C with rotation, and Dynabeads discarded. Cell lysates were then incubated with 5 µg of anti-FLAG antibody for 2 h at 4°C with rotation. 50 µl of Protein G Dynabeads were added, and incubated for 1 h at 4°C with rotation. Samples were washed four times with High Salt Buffer (50 mM Tris, 900 mM NaCl, 30 mM NaPyrophosphate, 50 mM NaF, 1% NP-40, 0.1% SDS), then eluted using NuPAGE 4x LDS sample buffer (Invitrogen) and heating at 70°C for 10 min. Samples were denatured and resolved by SDS-PAGE and western blotting as above.

For immunoprecipitation of recombinant MDA5 protein, MDA5 was diluted in MDA5 storage buffer (final concentration 0.0875 mg/ml). mAb 16 antibody-coupled Dynabeads were resuspended in MDA5 solution and incubated for 45 minutes at 25 °C. Beads were washed in Interaction buffer (25 mM HEPES-NaOH, 150 mM NaCl, 2.5 % glycerol, 0.1% (v/v) Tween-20, 1 mM DTT; pH 7.4 at 25 °C) supplemented with 1 mM CaCl_2_ as indicated in the corresponding figure. Beads were resuspended in ANXA2 diluted in Interaction buffer to a final concentration of 10 μM and incubated for 30 minutes at 25 °C. Beads were washed in Interaction buffer and samples were eluted by addition of 2 x Laemmli buffer and incubation for 10 minutes at 95 °C.

### SILAC mass spectrometry

THP1 cells were labelled with ‘light’ (R0K0) or ‘heavy’ (R10K8) SILAC RPMI medium (DC Biosciences) supplemented with 10% v/v SILAC Dyalised foetal bovine serum (DS1006, DC Biosciences) and 2 mM L-glutamine, and cryopreserved in multiple aliquots after 7 doublings, to allow complete incorporation of labelled isotopes. Cells were thawed and grown for an additional 4 days in respective labelled medium. 10 million THP1 cells were plated into 10 cm tissue culture dishes in the presence of 10 ng/ml PMA for 24 h. Heavy labelled cells were infected with EMCV at MOI = 10 for 20 hr, and light labelled cells were left uninfected. Cells were lysed and MDA5 was immunoprecipitated as described above, using a combined lysate of 500 µg infected and 500 µg uninfected lysate. Samples were eluted by incubating MDA5-bound Dynabeads with 200 mM glycine at pH 2.0 for 10 min. IP-eluates were subsequently processed for mass spectrometry by reduction/alkylation of cysteine and purification on SOLA C18 cartridges (ThermoFisher). Peptides were analysed on a LC/MS/MS platform consisting of Dionex Ultimate UHPLC and Q-Exactive mass spectrometer (both ThermoFisher) using a 60 minutes linear gradient from 2-35% Acetonitrile in 5% DMSO in 0.1% formic acid in a 50cm Easyspray column. Data was acquired in data dependent mode with a mass resolution of 70000 and a precursor mass range of 380-1800 M/z. The most abundant 15 precursor ions were selected for subsequent MS/MS analysis at a resolution of 17500 for up to 128ms. LC-MS/MS data were analysed in Fragpipe 23 (*64*) using SILAC standard parameters and further visualized with Perseus 2.1.5 (*65*) after normalisation and missing number imputation based on a downshifted normal distribution and plotted as volcano plot with permutation based FDR control. Full data can be found in Supplementary Data 1.

### Generation of EMCV-RNA and IAV-RNA

For generation of EMCV-RNA, HeLa cells were plated at 5x 10^6^ cells in 10 cm dishes and incubated overnight. Cell culture medium was exchanged, and cells invested with EMCV at MOI = 0.1 for 20 h. Cells were washed with PBS and RNA extracted with 1 ml of TRI Reagent solution (ThermoFisher, AM9738) according to manufacturer’s instructions. Purified RNA was treated with alkaline phosphatase (Roche 10713023001) according to manufacturer’s instructions, using 10 units of enzyme per 5 μg of RNA and incubating at 50°C for 1 h. RNA was purified by phenol:chloroform:iaa (Sigma), followed by ethanol precipitation. RNA was resuspended in RNase-free water and stored at −80°C in single-use aliquots. For generation of IAV-RNA, MDCK cells were infected with IAV HKx31 at MOI 5 for 8 h. RNA was extracted and purified as above, but without alkaline phosphatase treatment.

### Stimulation RNA ligands or virus infection

A549 cells were plated in 6-well plates at 2.5 x10^5^ cells/well and incubated overnight. RNA ligands were transfected in the amounts indicated in figures using Lipofectamine2000 (ThermoFisher) in OptiMEM (Gibco) according to manufacturer’s instructions. Cells were incubated for 16 h.

For IAV infection, cell culture medium was exchanged for a minimum volume of serum-free DMEM containing virus at the indicated MOI. Cells were incubated at 37°C for 1 h, with agitation every 20 min. Virus containing-medium was exchanged for fresh cell culture medium and cells incubated for indicated times. For EMCV and WNV infection, cell culture medium was exchanged for fresh medium containing virus at the indicated MOI, and cells incubated for indicated times.

### MDA5 oligomerisation assay

A549 cells were plated in 6-well plates at 3 x 10^5^ cells/well and incubated overnight. 800 ng/well of HeLa-EMCV-RNA was complexed with 5 µl of Lipofectamine2000 in OptiMEM according to manufacturer’s instructions and added to cells, and cells incubated for the indicated times. Cells were lifted using trypsin treatment to obtain a single-cell suspension, transferred to microcentrifuge tube, washed with PBS, and resuspended in 500 µl of disuccinimidyl suberate (DSS; ThermoFisher 21655) at 1 mM in conjugation buffer (100 mM sodium phosphate, 150 mM NaCl, 20 mM HEPES, 100 mM bicarbonate). Cells were rotated at room temperature for 30 min, prior to addition of 25 µl of 1 M Tris. Cells were rotated a further 15 min, and centrifuged at 300 x g. Cells were lysed and resolved by SDS-PAGE and Western blot as above, but without addition with 2-mercaptoethanol. MDA5 oligomers were detected by immunoblotting with mAb 17.

### RT-qPCR

RNA was extracted from cells using RNeasy Kit (Qiagen) according to manufacturer’s instructions, including DNaseI treatment to remove contaminating genomic DNA. RNA (1 μg) was converted into cDNA using SuperScript III Reverse Transcriptase Kit (Thermo Fisher Scientific) and random hexamer primers according to manufacturer’s instructions. cDNA was diluted to 100 ng/μl and quantitative PCR was performed using SYBR Green PCR Master Mix (Thermo Fisher Scientific) with gene-specific probes detailed in Supplementary Table 1. Assays were performed on QuantStudio 6 Flex Real-Time PCR machines (Thermo Fisher Scientific).

### RNA sequencing

For RNA extracted from THP1 cells associated with the EMCV infection experiment, libraries were generated using an in-house multiplex RNAseq method (MHTP, Hudson Medical Genomics Facility) as previously described (*66, 67*). Libraries were prepared using 25 ng of total RNA input. An 8 bp sample index and a 10 bp unique molecular identifier (UMI) were added during initial poly(A) priming and pooled samples were amplified using a template switching oligonucleotide. The Illumina P5 sequences were added by tagmentation by Nextera transposase and PCR. The library was designed so that the forward read (R1) sequenced the cDNA in the sense direction for transcript identification, while the reverse read (R2) sequences contained the index and then the 10 bp UMI. Paired-end sequencing was performed on a NextSeq2000 run using a P2 100cycle kit (cDNA reads generated 111 nt). Base calling was performed using Dragen BCLConvert (v3.10.11).

For RNA extracted from A549 cells and THP1 cells infected with WNV, a bulk RNA barcoding and sequencing (BRBseq) approach was used (*68*), with libraries generated using the MERCURIUS BRBseq kit (Mercurius, 1101) as per manufacturer’s instructions). Libraries were prepared using 500 ng of RNA input. Barcoded oligo(dT) primers containing 14 bp sample indexes and 14 bp UMI sequences were added during first-strand synthesis, with samples pooled for subsequent library preparation steps. Full-length cDNA was fragmented and tagged using Tn5 transposase, then a unique dual indexing strategy was used to add Illumina-compatible i5 and i7 sequences for PCR amplification. Pooled samples were split across two sequencing lanes and paired-end sequencing was performed on a NovaSeq X Plus (cDNA reads generated 150 nt), with base calling performed using DRAGEN. Raw sequencing reads from both sequencing lanes were combined for file processing steps, to enable accurate UMI deduplication.

### RNAseq analysis

RNA-seq read alignment and UMI counting was performed in R (https://www.r-project.org); v4.1.1, THP1 cells in EMCV experiment; v4.3.3, A549 cells and THP1 cells in WNV experiments). The scPipe (*69*) package (v1.14.0) was employed to process and de-multiplex the data. FASTQ files from BRB-seq remained multiplexed for initial stages of file processing; FASTQ files from in-house RNA sequencing had already been demultiplexed to obtain sample-specific files which contained the index and UMI sequences in the read header, but headers were reformatted using awk for compatibility with scPipe. A combined multiplexed FASTQ file was created from the R1 and R2 FASTQ files, by trimming the sample index and UMI sequences and storing them in the read header, using the sc_trim_barcode function (with bs2 = 0, bl2 = 8, us = 8, ul = 10 for in-house RNA-seq; bs2 = 0, bl2 = 14, us = 14, ul = 14 for BRB-seq). Read alignment was performed on the combined multiplexed FASTQ file using the RSubread (*70*) package (v2.6.4). An index was built using the Ensembl *Homo sapiens* GRCh38 primary assembly genome file and alignment was performed with default settings. Aligned reads were mapped to exons using the sc_exon_mapping function with an Ensembl *Homo sapiens* GRCh38 GFF3 genome annotation file (v110, THP1 cells in EMCV experiment; v113, A549 and THP1 cells in WNV experiments). Reads mapping to exons were associated with each individual sample using the sc_demultiplex function, taking the UMI into account, and an overall count for each gene for each was sample was generated using the sc_gene_counting function (with UMI_cor = 1). Subsequent analyses were performed in R (v4.1.2, THP1 cells in EMCV experiment; v4.4.2, A549 and THP1 cells in WNV experiments).

Additional gene annotation was obtained using the biomaRt (*71*) package (v2.50.1, THP1 cells in EMCV experiment; v2.62.1, A549 and THP1 cells in WNV experiments) and a DGEList object was created with the counts and gene annotation using the edgeR (*72*) package (v3.36.0, THP1 cells in EMCV experiment; v4.4.2, A549 and THP1 cells in WNV experiments). A design matrix was constructed incorporating the treatment groups and experimental replicate. Lowly expressed genes were removed using the filterByExpr function and normalisation factors were calculated using the TMM method (*73*). Counts were transformed using the voom method (*74*) and a linear model was fit using the edgeR voomLmFit function. Differential gene expression analyses were performed using the limma (*75*) (v3.50.0, THP1 cells in EMCV experiment; v3.62.2, A549 cells; v3.65.4, THP1 cells in WNV experiments) package. Groups were compared using the contrasts.fit function and moderated t-statistics were calculated without a fold-change threshold using the eBayes function (*76*) (A549 and THP1 cells in WNV experiments) or were calculated using the treat function (*77*) with a 1.2-fold cut-off (THP1 cells in EMCV experiment). Differentially expressed genes were determined using a false discovery rate (FDR) adjusted *p*-value < 0.05. Heat maps were made using the pheatmap package (https://CRAN.R-project.org/package=pheatmap; v1.0.12). Hallmark gene sets were obtained from the Broad Institute Molecular Signature Database (*78*), via the msigdbr (*79*) package (v7.4.1, THP1 cells in EMCV experiment; v24.1.0, A549 and THP1 cells in WNV experiments). Log_2_ counts per million (CPM) expression values were calculated using the edgeR cpm function. Relative log_2_ CPM expression values were calculated for each gene by subtracting the average log_2_ CPM value of untreated wild type controls. Where relevant, log_2_ fold changes between different groups were also shown, indicating adjusted *p*-values as * *p* < 0.05, ** *p* < 0.01 and *** *p* < 0.001. Gene set tests were performed using the cameraPR function (*80*) from the limma package on moderated *t*-statistics recalculated without a fold-change threshold, using the eBayes function (*76*). Test results were presented as heat maps showing the average log_2_ fold changes of genes in each set (rows) for relevant comparisons (columns), with scales truncated at ±2. The significance of each gene set was indicated by the text as the −log_10_ FDR-adjusted p-value threshold, with asterisks denoting p < 0.05, 2 denoting p < 0.01, 3 denoting p < 0.001 and so forth.

### iPSC-derived macrophages

Kolf2 iPSCs were routinely cultured on Matrigel-coated 6-well plates in mTeSR Plus medium (STEMCELL Technologies) and verified for normal karyotype and absence of mycoplasma. Cells were passaged at 70–80% confluency using 0.5 mM EDTA (Gibco). To generate ANXA2-KO cells, Alt-R CRISPR-Cas9 guide RNAs (Integrated DNA Technologies) were used according to manufacturer’s instructions. Briefly, crRNA targeting ANXA2 (Hs.Cas9.ANXA2.1.AA) was complexed with tracrRNA ATTO 550 (Integrated DNA Technologies) and TrueCut Cas9 Protein (A36498, ThermoFisher), then introduced into cells using the Lonza 4D Nucleofector and P3 Primary Cell 4D X Kit (V4XP-3032, Lonza). Single-cell clones were screened for homozygous disruption of target genes by sequencing.

To generate macrophages, iPSCs were differentiated under a modified serum-free monolayer protocol adapted from H. Hosseini Far et al (in preparation). Briefly, iPSC with 70– 80% confluency was dissociated and seeded at a density of 1.0–1.5 x 10^4^ cells/cm^2^ with mTeSR Plus medium supplemented with 10 μM Y-27632 (ROCK inhibitor; STEMCELL Technologies). The next day, differentiation was initiated using Essential 6 (E6) medium (ThermoFisher) supplemented with 0.1% polyvinyl alcohol (Sigma) and methyl cellulose (Sigma), along with specific combinations of growth factors and small molecules added sequentially to guide lineage commitment. On **Day 1**, cells were treated with 10 ng/ml Activin A, 10 ng/ml BMP4 (R&D System), 20 ng/ml FGF2 (Peprotech), 6 µM CHIR99021 (Tocris), and 100 nM PIK90 (Sigma). On **Day 2**, differentiation was directed toward a mesendothelial fate by adding 10 ng/ml BMP4, 20 ng/ml FGF2, 50 ng/ml VEGF (Peprotech), and 1 µM C59 (Tocris). On **Day 3**, the cells were treated with 20 ng/ml BMP4, 20 ng/ml FGF2, 50 ng/ml VEGF, 50 ng/ml SCF (Peprotech), and 1 µM C59, promoting hemogenic endothelium development. From **Day 4 to Day 9**, cells were maintained in E6 medium supplemented with 20 ng/ml FGF2, 50 ng/ml VEGF, 50 ng/ml SCF, and 50 ng/ml IL-3 (Peprotech) to support hematopoietic specification and expansion of early progenitors. Floating hematopoietic progenitors appeared by Day 7 and were harvested from Day 9 onward. The floating progenitor cells were cultured in mCSF (Peprotech)-containing E6 media for 3 weeks to mature into macrophages.

For analysis of cell surface markers, cells were washed twice in FACS buffer (2% FCS in PBS) and surface stained with CD34-PE, CD14-BV421, CD163-APC, CD11b-APC-Cy7, and CD16-PE-Cy7 for 30 min at 4°C. All antibodies were diluted 1:50 in FACS buffer. Cells were washed twice in FACS buffer and acquired on a Fortessa flow cytometer (BD Biosciences).

### iPSC-derived neuronal progenitor cells

Kolf2 iPSCs were maintained as above. For neural induction, iPSCs were dissociated into single cells using Accutase (STEMCELL Technologies) and seeded at a density of 1 × 10⁶ cells/mL onto fresh Matrigel-coated plates in StemDiff Neural Induction Medium (STEMCELL Technologies) supplemented with the SMAD Inhibition Supplement (STEMCELL Technologies) and 10 μM Y-27632. Cells were maintained in Neural Induction Medium with SMAD Inhibition Supplement throughout the induction phase, with daily medium changes, and were passaged using Accutase every 5–7 days at the same seeding density with 10 μM Y-27632. After the third passage, cells exhibiting neural rosette morphology were dissociated and seeded at a density of 2 × 10⁵ cells/cm² onto plates pre-coated with poly-L-ornithine (15 μg/ml) and laminin (10 μg/ml). Culture medium was then switched to StemDiff Neural Progenitor Medium (STEMCELL Technologies) supplemented with the manufacturer-recommended supplements to support neural progenitor cell expansion. Neural progenitor cells were maintained in Neural Progenitor Medium and expanded for downstream applications. Cells were plated in 12-well dishes at 2.5 x 10^5^ cells/well for WNV infection experiments.

### Cytometric bead array

Cytokines were measured in cell culture supernatants using BD Cytometric Bead Array Flex kits for human IL-6, IL-8, CXCL10, IL-10, IFNα, TNFα, and CCL5. Capture beads from each Flex kit were vortexed before use and 0.1μl/well of each bead was diluted in assay buffer (0.1% BSA, 2 mM EDTA in PBS), for a total volume of 5 μl per well. Detection reagent was also diluted in assay buffer using 0.1μl/well of each PE-conjugated detection antibody in total volume of 5 μl per well. Serially diluted cytokine standards and neat supernatant samples were added to a V-bottom plate at 5 μl per well, followed by 5 μl per well of the capture beads and 5 μl per well of the detection reagent. Samples were incubated in the dark at room temperature. After 2 hours, 200 μl of assay buffer was added to the wells and the plate was centrifuged at 1000 g for 5 mins. Following centrifugation, liquid was carefully removed from the wells and the pelleted beads were resuspended in 200 μl of 2% PFA in PBS. The plate was incubated for a further 30 minutes at room temperature in the dark to ensure inactivation of any viral particles. The plate was centrifuged again and the beads resuspended in 100 μl assay buffer for acquisition on a Cytek Aurora in Conventional Mode.

Data was analysed in FlowJo v10 (BD Biosciences), with each cytokine bead population identified via its position based on fluorescence in the APC and APC-Cy7 channels. The Mean Fluorescence Intensity (MFI) of the PE channel was then measured for each population of beads in each standard and sample. To determine cytokine concentrations, log_10_(concentration) values were plotted against the measured MFIs for the standards to generate a standard curve for each cytokine, which was then used to interpolate the concentrations of cytokines in the supernatant samples.

## Acknowledgments

We acknowledge the Hudson Genomics at the Hudson Institute of Medical Research for technical support with sequencing experiments. This research was supported by the Scientific Service Units (SSU) of Institute of Science and Technology Austria (ISTA) through resources provided by the Lab Support Facility (LSF) Institute.

## Author contributions

N.G.S and J.R conceptualized the project; N.G.S, P.H., C.B. and S.D. supervised the project; N.G.S, G.W-M., D.M., R.L., N.C., S.S, H.H-F., A.M. and R.M. conducted experiments; M.T. provided essential reagents; N.G.S, A.McA., J.G. and R.M. analysed the data; and N.G.S, prepared the manuscript. All authors read and approved the manuscript.

## Data and materials availability

All data are available in the main text or the supplementary materials. All requests for reagents, resources, or further information should be addressed to and will be fulfilled by the lead contact, Natalia G. Sampaio. Raw bulk RNA sequencing data are deposited in the NCBI Gene Expression Omnibus: GSE301029 and GSE301031. This paper does not report original code.

## References

1. A. G. Dias Junior, N. G. Sampaio, J. Rehwinkel, A Balancing Act: MDA5 in Antiviral Immunity and Autoinflammation. Trends in Microbiology 27, 75–85 (2018).

2. J. Rehwinkel, M. U. Gack, RIG-I-like receptors: their regulation and roles in RNA sensing. Nature Reviews Immunology 20, 537–551 (2020).

3. B. Wu, A. Peisley, C. Richards, H. Yao, X. Zeng, C. Lin, F. Chu, T. Walz, S. Hur, Structural basis for dsRNA recognition, filament formation, and antiviral signal activation by MDA5. Cell 152, 276–289 (2013).

4. I. T. Lamborn, H. Jing, Y. Zhang, S. B. Drutman, J. K. Abbott, S. Munir, S. Bade, H. M. Murdock, C. P. Santos, L. G. Brock, E. Masutani, E. Y. Fordjour, J. J. McElwee, J. D. Hughes, D. P. Nichols, A. Belkadi, A. J. Oler, C. S. Happel, H. F. Matthews, L. Abel, P. L. Collins, K. Subbarao, E. W. Gelfand, M. J. Ciancanelli, J.-L. Casanova, H. C. Su, Recurrent rhinovirus infections in a child with inherited MDA5 deficiency. The Journal of Experimental Medicine 214, 1949 - 1972 (2017).

5. S. Asgari, L. J. Schlapbach, S. Anchisi, C. Hammer, I. Bartha, T. Junier, G. Mottet-Osman, K. M. Posfay-Barbe, D. Longchamp, M. Stocker, S. Cordey, L. Kaiser, T. Riedel, T. Kenna, D. Long, A. Schibler, A. Telenti, C. Tapparel, P. J. McLaren, D. Garcin, J. Fellay, Severe viral respiratory infections in children with IFIH1 loss-of-function mutations. Proceedings of the National Academy of Sciences 114, 8342–8347 (2017).

6. S. Nejentsev, N. Walker, D. Riches, M. Egholm, J. A. Todd, Rare variants of IFIH1, a gene implicated in antiviral responses, protect against type 1 diabetes. Science (New York, N.Y.) 324, 387–389 (2009).

7. D. J. Smyth, J. D. Cooper, R. Bailey, S. Field, O. Burren, L. J. Smink, C. Guja, C. Ionescu-Tirgoviste, B. Widmer, D. B. Dunger, D. A. Savage, N. M. Walker, D. G. Clayton, J. A. Todd, A genome-wide association study of nonsynonymous SNPs identifies a type 1 diabetes locus in the interferon-induced helicase (IFIH1) region. Nature Genetics 38, 617–619 (2006).

8. R. Singh, J. D. Joiner, A. Herrero Del Valle, M. Zwaagstra, I. Jobe, B. J. Ferguson, F. J. M. van Kuppeveld, Y. Modis, Molecular basis of autoimmune disease protection by MDA5 variants. Cell Rep 44, 115754 (2025).

9. M. Funabiki, H. Kato, Y. Miyachi, H. Toki, H. Motegi, M. Inoue, O. Minowa, A. Yoshida, K. Deguchi, H. Sato, S. Ito, T. Shiroishi, K. Takeyasu, T. Noda, T. Fujita, Autoimmune disorders associated with gain of function of the intracellular sensor MDA5. Immunity 40, 199–212 (2014).

10. G. I. Rice, S. Park, F. Gavazzi, L. A. Adang, L. A. Ayuk, L. Van Eyck, L. Seabra, C. Barrea, R. Battini, A. Belot, S. Berg, T. Billette de Villemeur, A. E. Bley, L. Blumkin, O. Boespflug-Tanguy, T. A. Briggs, E. Brimble, R. C. Dale, N. Darin, F. G. Debray, V. De Giorgis, J. Denecke, D. Doummar, G. Drake Af Hagelsrum, D. Eleftheriou, M. Estienne, E. Fazzi, F. Feillet, J. Galli, N. Hartog, J. Harvengt, B. Heron, D. Heron, D. A. Kelly, D. Lev, V. Levrat, J. H. Livingston, I. Marti, C. Mignot, F. Mochel, M. C. Nougues, I. Oppermann, B. Pérez-Dueñas, B. Popp, M. P. Rodero, D. Rodriguez, V. Saletti, C. Sharpe, D. Tonduti, G. Vadlamani, K. Van Haren, M. Tomas Vila, J. Vogt, E. Wassmer, A. Wiedemann, C. J. Wilson, A. Zerem, C. Zweier, S. M. Zuberi, S. Orcesi, A. L. Vanderver, S. Hur, Y. J. Crow, Genetic and phenotypic spectrum associated with IFIH1 gain-of-function. Hum Mutat 41, 837–849 (2020).

11. D. Blanco-Melo, B. E. Nilsson-Payant, W. C. Liu, S. Uhl, D. Hoagland, R. Møller, T. X. Jordan, K. Oishi, M. Panis, D. Sachs, T. T. Wang, R. E. Schwartz, J. K. Lim, R. A. Albrecht, B. R. tenOever, Imbalanced Host Response to SARS-CoV-2 Drives Development of COVID-19. Cell 181, 1036–1045.e1039 (2020).

12. R. Karki, B. R. Sharma, S. Tuladhar, E. P. Williams, L. Zalduondo, P. Samir, M. Zheng, B. Sundaram, B. Banoth, R. K. S. Malireddi, P. Schreiner, G. Neale, P. Vogel, R. Webby, C. B. Jonsson, T. D. Kanneganti, Synergism of TNF-α and IFN-γ Triggers Inflammatory Cell Death, Tissue Damage, and Mortality in SARS-CoV-2 Infection and Cytokine Shock Syndromes. Cell 184, 149–168.e117 (2021).

13. A. Peisley, C. Lin, B. Wu, M. Orme-Johnson, M. Liu, T. Walz, S. Hur, Cooperative assembly and dynamic disassembly of MDA5 filaments for viral dsRNA recognition. Proceedings of the National Academy of Sciences 108, 21010–21015 (2011).

14. F. Hou, L. Sun, H. Zheng, B. Skaug, Q. X. Jiang, Z. J. Chen, MAVS forms functional prion-like aggregates to activate and propagate antiviral innate immune response. Cell 146, 448–461 (2011).

15. D. Acharya, R. Reis, M. Volcic, G. Liu, M. K. Wang, B. S. Chia, R. Nchioua, R. Groß, J. Münch, F. Kirchhoff, K. M. J. Sparrer, M. U. Gack, Actin cytoskeleton remodeling primes RIG-I-like receptor activation. Cell 185, 3588–3602.e3521 (2022).

16. X. Lang, T. Tang, T. Jin, C. Ding, R. Zhou, W. Jiang, TRIM65-catalized ubiquitination is essential for MDA5-mediated antiviral innate immunity. The Journal of experimental medicine 214, 459–473 (2017).

17. M.-M. Hu, C.-Y. Liao, Q. Yang, X.-Q. Xie, H.-B. Shu, Innate immunity to RNA virus is regulated by temporal and reversible sumoylation of RIG-I and MDA5. The Journal of experimental medicine, jem.20161015 (2017).

18. L. Sarkar, G. Liu, D. Acharya, J. Zhu, Z. Sayyad, M. U. Gack, MDA5 ISGylation is crucial for immune signaling to control viral replication and pathogenesis. Proc Natl Acad Sci U S A 122, e2420190122 (2025).

19. G. Liu, J.-H. Lee, Z. M. Parker, D. Acharya, J. J. Chiang, M. v. Gent, W. Riedl, M. E. Davis-Gardner, E. Wies, C. Chiang, M. U. Gack, ISG15-dependent activation of the sensor MDA5 is antagonized by the SARS-CoV-2 papain-like protease to evade host innate immunity. Nature Microbiology 6, 467–478 (2021).

20. G. Li, J. Zhang, Z. Zhao, J. Wang, J. Li, W. Xu, Z. Cui, P. Sun, H. Yuan, T. Wang, K. Li, X. Bai, X. Ma, P. Li, Y. Fu, Y. Cao, H. Bao, D. Li, Z. Liu, N. Zhu, L. Tang, Z. Lu, RNF144B negatively regulates antiviral immunity by targeting MDA5 for autophagic degradation. Embo Rep 25, 4594–4624 (2024).

21. M. He, Z. Yang, L. Xie, J. Chen, S. Liu, L. Lu, Z. Li, B. Zheng, Y. Ye, Y. Lin, L. Bu, J. Xiao, Y. Zhong, P. Jia, Q. Li, Y. Liang, D. Guo, C. M. Li, P. Hou, RNF167 mediates atypical ubiquitylation and degradation of RLRs via two distinct proteolytic pathways. Nat Commun 16, 1920 (2025).

22. V. Rivero, J. Carrión-Cruz, D. López-García, M. L. DeDiego, The IFN-induced protein IFI27 binds MDA5 and counteracts its activation after SARS-CoV-2 infection. Front Cell Infect Microbiol 14, 1470924 (2024).

23. A. G. Harrison, D. Yang, J. G. Cahoon, T. Geng, Z. Cao, T. A. Karginov, Y. Hu, X. Li, C. C. Chiari, Y. Qyang, A. T. Vella, Z. Fan, S. K. Vanaja, V. A. Rathinam, C. A. Witczak, J. S. Bogan, P. Wang, UBXN9 governs GLUT4-mediated spatial confinement of RIG-I-like receptors and signaling. Nat Immunol 25, 2234–2246 (2024).

24. J.-P. Lin, Y.-K. Fan, H. M. Liu, The 14-3-3η chaperone protein promotes antiviral innate immunity via facilitating MDA5 oligomerization and intracellular redistribution. PLoS pathogens 15, e1007582 (2019).

25. W. Riedl, D. Acharya, J.-H. Lee, G. Liu, T. Serman, C. Chiang, Y. K. Chan, M. S. Diamond, M. U. Gack, Zika Virus NS3 Mimics a Cellular 14-3-3-Binding Motif to Antagonize RIG-I- and MDA5-Mediated Innate Immunity. Cell host & microbe 26, 493–503.e496 (2019).

26. P.-Y. Lui, L.-Y. R. Wong, T.-H. Ho, S. W. N. Au, C.-P. Chan, K.-H. Kok, D.-Y. Jin, PACT Facilitates RNA-Induced Activation of MDA5 by Promoting MDA5 Oligomerization. Journal of immunology (Baltimore, Md.: 1950) 199, 1846–1855 (2017).

27. L. E. Bazzone, J. Zhu, M. King, G. Liu, Z. Guo, C. R. MacKay, P. P. Kyawe, N. Qaisar, J. Rojas-Quintero, C. A. Owen, A. L. Brass, W. McDougall, C. E. Baer, T. Cashman, C. M. Trivedi, M. U. Gack, R. W. Finberg, E. A. Kurt-Jones, ADAM9 promotes type I interferon-mediated innate immunity during encephalomyocarditis virus infection. Nature Communications 15, 4153 (2024).

28. M. Shi, T. Jiang, M. Zhang, Q. Li, K. Liu, N. Lin, X. Wang, A. Jiang, Y. Gao, Y. Wang, S. Liu, L. Zhang, D. Li, P. Gao, Nucleic-acid-induced ZCCHC3 condensation promotes broad innate immune responses. Mol Cell 85, 962–975.e967 (2025).

29. S. Ahmad, X. Mu, F. Yang, E. Greenwald, J. W. Park, E. Jacob, C.-Z. Zhang, S. Hur, Breaching Self-Tolerance to Alu Duplex RNA Underlies MDA5-Mediated Inflammation. Cell 172, 797–810 (2018).

30. H. Kato, O. Takeuchi, S. Sato, M. Yoneyama, M. Yamamoto, K. Matsui, S. Uematsu, A. Jung, T. Kawai, K. J. Ishii, O. Yamaguchi, K. Otsu, T. Tsujimura, C.-S. Koh, C. R. E. Sousa, Y. Matsuura, T. Fujita, S. Akira, Differential roles of MDA5 and RIG-I helicases in the recognition of RNA viruses. Nature 441, 101–105 (2006).

31. V. Gerke, F. N. E. Gavins, M. Geisow, T. Grewal, J. K. Jaiswal, J. Nylandsted, U. Rescher, Annexins-a family of proteins with distinctive tastes for cell signaling and membrane dynamics. Nat Commun 15, 1574 (2024).

32. U. Rescher, D. Ruhe, C. Ludwig, N. Zobiack, V. Gerke, Annexin 2 is a phosphatidylinositol (4,5)-bisphosphate binding protein recruited to actin assembly sites at cellular membranes. J Cell Sci 117, 3473–3480 (2004).

33. N. R. Filipenko, T. J. MacLeod, C. S. Yoon, D. M. Waisman, Annexin A2 is a novel RNA-binding protein. J Biol Chem 279, 8723–8731 (2004).

34. Y. Liu, H. K. Myrvang, L. V. Dekker, Annexin A2 complexes with S100 proteins: structure, function and pharmacological manipulation. British Journal of Pharmacology 172, 1664–1676 (2015).

35. M. Jost, V. Gerke, Mapping of a regulatory important site for protein kinase C phosphorylation in the N-terminal domain of annexin II. Biochimica et Biophysica Acta (BBA) - Molecular Cell Research 1313, 283–289 (1996).

36. M. Kumar, M. Gouw, S. Michael, H. Sámano-Sánchez, R. Pancsa, J. Glavina, A. Diakogianni, J. A. Valverde, D. Bukirova, J. Čalyševa, N. Palopoli, N. E. Davey, L. B. Chemes, T. J. Gibson, ELM—the eukaryotic linear motif resource in 2020. Nucleic Acids Research 48, D296–D306 (2019).

37. S. Straub, N. G. Sampaio, Activation of cytosolic RNA sensors by endogenous ligands: roles in disease pathogenesis. Frontiers in Immunology 14, (2023).

38. J. S. Errett, M. S. Suthar, A. McMillan, M. S. Diamond, M. Gale, Jr., The essential, nonredundant roles of RIG-I and MDA5 in detecting and controlling West Nile virus infection. Journal of virology 87, 11416–11425 (2013).

39. A. E. L. Stone, R. Green, C. Wilkins, E. A. Hemann, M. Gale, RIG-I-like receptors direct inflammatory macrophage polarization against West Nile virus infection. Nature Communications 10, 3649–3616 (2019).

40. E. Genoyer, J. Wilson, J. M. Ames, C. Stokes, D. Moreno, N. Etzyon, A. Oberst, M. Gale, Jr., Exposure of negative-sense viral RNA in the cytoplasm initiates innate immunity to West Nile virus. Mol Cell 85, 1147–1161.e1149 (2025).

41. P. E. Ceccaldi, M. Lucas, P. Despres, New insights on the neuropathology of West Nile virus. FEMS Microbiol Lett 233, 1–6 (2004).

42. J. Rehwinkel, C. P. Tan, D. Goubau, O. Schulz, A. Pichlmair, K. Bier, N. Robb, F. Vreede, W. Barclay, E. Fodor, C. R. E. Sousa, RIG-I detects viral genomic RNA during negative-strand RNA virus infection. Cell 140, 397–408 (2010).

43. A. García-Sastre, A. Egorov, D. Matassov, S. Brandt, D. E. Levy, J. E. Durbin, P. Palese, T. Muster, Influenza A virus lacking the NS1 gene replicates in interferon-deficient systems. Virology 252, 324–330 (1998).

44. M. D. Park, A. Silvin, F. Ginhoux, M. Merad, Macrophages in health and disease. Cell 185, 4259–4279 (2022).

45. C. David, C. Verney, M. Si-Tahar, A. Guillon, The deadly dance of alveolar macrophages and influenza virus. Eur Respir Rev 33, (2024).

46. D. R. Getts, R. L. Terry, M. T. Getts, M. Müller, S. Rana, B. Shrestha, J. Radford, N. Van Rooijen, I. L. Campbell, N. J. King, Ly6c+ “inflammatory monocytes” are microglial precursors recruited in a pathogenic manner in West Nile virus encephalitis. J Exp Med 205, 2319–2337 (2008).

47. V. Gerke, C. E. Creutz, S. E. Moss, Annexins: linking Ca2+ signalling to membrane dynamics. Nature Reviews Molecular Cell Biology 6, 449–461 (2005).

48. E. Strand, H. Hollås, S. A. Sakya, S. Romanyuk, M. E. V. Saraste, A. K. Grindheim, S. S. Patil, A. Vedeler, Annexin A2 binds the internal ribosomal entry site of c-myc mRNA and regulates its translation. RNA Biol 18, 337–354 (2021).

49. A. Irurzun, J. Arroyo, A. Alvarez, L. Carrasco, Enhanced intracellular calcium concentration during poliovirus infection. J Virol 69, 5142–5146 (1995).

50. D. Máthé, G. Szalay, L. Cseri, Z. Kis, B. Pályi, G. Földes, N. Kovács, A. Fülöp, Á. Szepesi, P. Hajdrik, A. Csomos, Á. Zsembery, K. Kádár, G. Katona, Z. Mucsi, B. J. Rózsa, E. Kovács, Monitoring correlates of SARS-CoV-2 infection in cell culture using a two-photon-active calcium-sensitive dye. Cellular & Molecular Biology Letters 29, 105 (2024).

51. J.-P. Brunet, J. Cotte-Laffitte, C. Linxe, A.-M. Quero, M. Géniteau-Legendre, A. Servin, Rotavirus Infection Induces an Increase in Intracellular Calcium Concentration in Human Intestinal Epithelial Cells: Role in Microvillar Actin Alteration. Journal of Virology 74, 2323–2332 (2000).

52. J. T. Gebert, F. J. Scribano, K. A. Engevik, E. M. Huleatt, M. R. Eledge, L. E. Dorn, A. A. Philip, T. Kawagishi, H. B. Greenberg, J. T. Patton, J. M. Hyser, Viroporin activity is necessary for intercellular calcium signals that contribute to viral pathogenesis. Sci Adv 11, eadq8115 (2025).

53. S. V. Scherbik, M. A. Brinton, Virus-induced Ca2+ influx extends survival of west nile virus-infected cells. J Virol 84, 8721–8731 (2010).

54. H. Lian, R. Zang, J. Wei, W. Ye, M.-M. Hu, Y.-D. Chen, X.-N. Zhang, Y. Guo, C.-Q. Lei, Q. Yang, W.-W. Luo, S. Li, H.-B. Shu, The Zinc-Finger Protein ZCCHC3 Binds RNA and Facilitates Viral RNA Sensing and Activation of the RIG-I-like Receptors. Immunity 49, 438–448.e435 (2018).

55. Y. C. Liao, M. S. Fernandopulle, G. Wang, H. Choi, L. Hao, C. M. Drerup, R. Patel, S. Qamar, J. Nixon-Abell, Y. Shen, W. Meadows, M. Vendruscolo, T. P. J. Knowles, M. Nelson, M. A. Czekalska, G. Musteikyte, M. A. Gachechiladze, C. A. Stephens, H. A. Pasolli, L. R. Forrest, P. St George-Hyslop, J. Lippincott-Schwartz, M. E. Ward, RNA Granules Hitchhike on Lysosomes for Long-Distance Transport, Using Annexin A11 as a Molecular Tether. Cell 179, 147–164.e120 (2019).

56. W. W. Yim, H. Yamamoto, N. Mizushima, Annexins A1 and A2 are recruited to larger lysosomal injuries independently of ESCRTs to promote repair. FEBS Lett 596, 991–1003 (2022).

57. J. R. Taylor, J. G. Skeate, W. M. Kast, Annexin A2 in Virus Infection. Frontiers in microbiology 9, 2954 (2018).

58. J. Chen, Y. Liu, S. Xia, X. Ye, L. Chen, Annexin A2 (ANXA2) regulates the transcription and alternative splicing of inflammatory genes in renal tubular epithelial cells. Bmc Genomics 23, 544 (2022).

59. A. Basters, K. P. Knobeloch, G. Fritz, USP18 - a multifunctional component in the interferon response. Biosci Rep 38, (2018).

60. A. Yoshimura, M. Ito, S. Chikuma, T. Akanuma, H. Nakatsukasa, Negative Regulation of Cytokine Signaling in Immunity. Cold Spring Harb Perspect Biol 10, (2018).

61. J. Hertzog, A. G. Dias Junior, R. E. Rigby, C. L. Donald, A. Mayer, E. Sezgin, C. Song, B. Jin, P. Hublitz, C. Eggeling, A. Kohl, J. Rehwinkel, Infection with a Brazilian isolate of Zika virus generates RIG-I stimulatory RNA and the viral NS5 protein blocks type I IFN induction and signaling. Eur J Immunol 48, 1120–1136 (2018).

62. S. T. Sarvestani, M. D. Tate, J. M. Moffat, A. M. Jacobi, M. A. Behlke, A. R. Miller, S. A. Beckham, C. E. McCoy, W. Chen, J. D. Mintern, M. O’Keeffe, M. John, B. R. Williams, M. P. Gantier, Inosine-mediated modulation of RNA sensing by Toll-like receptor 7 (TLR7) and TLR8. J Virol 88, 799–810 (2014).

63. S. D. Gradia, J. P. Ishida, M. S. Tsai, C. Jeans, J. A. Tainer, J. O. Fuss, MacroBac: New Technologies for Robust and Efficient Large-Scale Production of Recombinant Multiprotein Complexes. Methods Enzymol 592, 1–26 (2017).

64. A. T. Kong, F. V. Leprevost, D. M. Avtonomov, D. Mellacheruvu, A. I. Nesvizhskii, MSFragger: ultrafast and comprehensive peptide identification in mass spectrometry–based proteomics. Nature Methods 14, 513–520 (2017).

65. S. Tyanova, T. Temu, P. Sinitcyn, A. Carlson, M. Y. Hein, T. Geiger, M. Mann, J. Cox, The Perseus computational platform for comprehensive analysis of (prote)omics data. Nature Methods 13, 731–740 (2016).

66. A. Grubman, X. Y. Choo, G. Chew, J. F. Ouyang, G. Sun, N. P. Croft, F. J. Rossello, R. Simmons, S. Buckberry, D. V. Landin, J. Pflueger, T. H. Vandekolk, Z. Abay, Y. Zhou, X. Liu, J. Chen, M. Larcombe, J. M. Haynes, C. McLean, S. Williams, S. Y. Chai, T. Wilson, R. Lister, C. W. Pouton, A. W. Purcell, O. J. L. Rackham, E. Petretto, J. M. Polo, Transcriptional signature in microglia associated with Aβ plaque phagocytosis. Nature Communications 12, 3015 (2021).

67. G. L. D’Adamo, M. Chonwerawong, L. J. Gearing, V. R. Marcelino, J. A. Gould, E. L. Rutten, S. M. Solari, P. W. R. Khoo, T. J. Wilson, T. Thomason, E. L. Gulliver, P. J. Hertzog, E. M. Giles, S. C. Forster, Bacterial clade-specific analysis identifies distinct epithelial responses in inflammatory bowel disease. Cell Rep Med 4, 101124 (2023).

68. D. Alpern, V. Gardeux, J. Russeil, B. Mangeat, A. C. A. Meireles-Filho, R. Breysse, D. Hacker, B. Deplancke, BRB-seq: ultra-affordable high-throughput transcriptomics enabled by bulk RNA barcoding and sequencing. Genome Biology 20, 71 (2019).

69. L. Tian, S. Su, X. Dong, D. Amann-Zalcenstein, C. Biben, A. Seidi, D. J. Hilton, S. H. Naik, M. E. Ritchie, scPipe: A flexible R/Bioconductor preprocessing pipeline for single-cell RNA-sequencing data. PLoS Comput Biol 14, e1006361 (2018).

70. Y. Liao, G. K. Smyth, W. Shi, The R package Rsubread is easier, faster, cheaper and better for alignment and quantification of RNA sequencing reads. Nucleic Acids Res 47, e47 (2019).

71. S. Durinck, P. T. Spellman, E. Birney, W. Huber, Mapping identifiers for the integration of genomic datasets with the R/Bioconductor package biomaRt. Nat Protoc 4, 1184–1191 (2009).

72. M. D. Robinson, D. J. McCarthy, G. K. Smyth, edgeR: a Bioconductor package for differential expression analysis of digital gene expression data. Bioinformatics 26, 139–140 (2010).

73. M. D. Robinson, A. Oshlack, A scaling normalization method for differential expression analysis of RNA-seq data. Genome Biol 11, R25 (2010).

74. C. W. Law, Y. Chen, W. Shi, G. K. Smyth, voom: Precision weights unlock linear model analysis tools for RNA-seq read counts. Genome Biol 15, R29 (2014).

75. M. E. Ritchie, B. Phipson, D. Wu, Y. Hu, C. W. Law, W. Shi, G. K. Smyth, limma powers differential expression analyses for RNA-sequencing and microarray studies. Nucleic Acids Res 43, e47 (2015).

76. B. Phipson, S. Lee, I. J. Majewski, W. S. Alexander, G. K. Smyth, Robust hyperparameter estimation protects against hypervariable genes and improves power to detect differential expression. The Annals of Applied Statistics 10, 946–963, 918 (2016).

77. D. J. McCarthy, G. K. Smyth, Testing significance relative to a fold-change threshold is a TREAT. Bioinformatics 25, 765–771 (2009).

78. A. Subramanian, P. Tamayo, V. K. Mootha, S. Mukherjee, B. L. Ebert, M. A. Gillette, A. Paulovich, S. L. Pomeroy, T. R. Golub, E. S. Lander, J. P. Mesirov, Gene set enrichment analysis: a knowledge-based approach for interpreting genome-wide expression profiles. Proc Natl Acad Sci U S A 102, 15545–15550 (2005).

79. A. Liberzon, C. Birger, H. Thorvaldsdóttir, M. Ghandi, J. P. Mesirov, P. Tamayo, The Molecular Signatures Database (MSigDB) hallmark gene set collection. Cell Syst 1, 417–425 (2015).

80. D. Wu, G. K. Smyth, Camera: a competitive gene set test accounting for inter-gene correlation. Nucleic Acids Res 40, e133 (2012).

