## Supplemental material for "Annexin A2 mediates RIG-I-like receptor responses to viral infection"

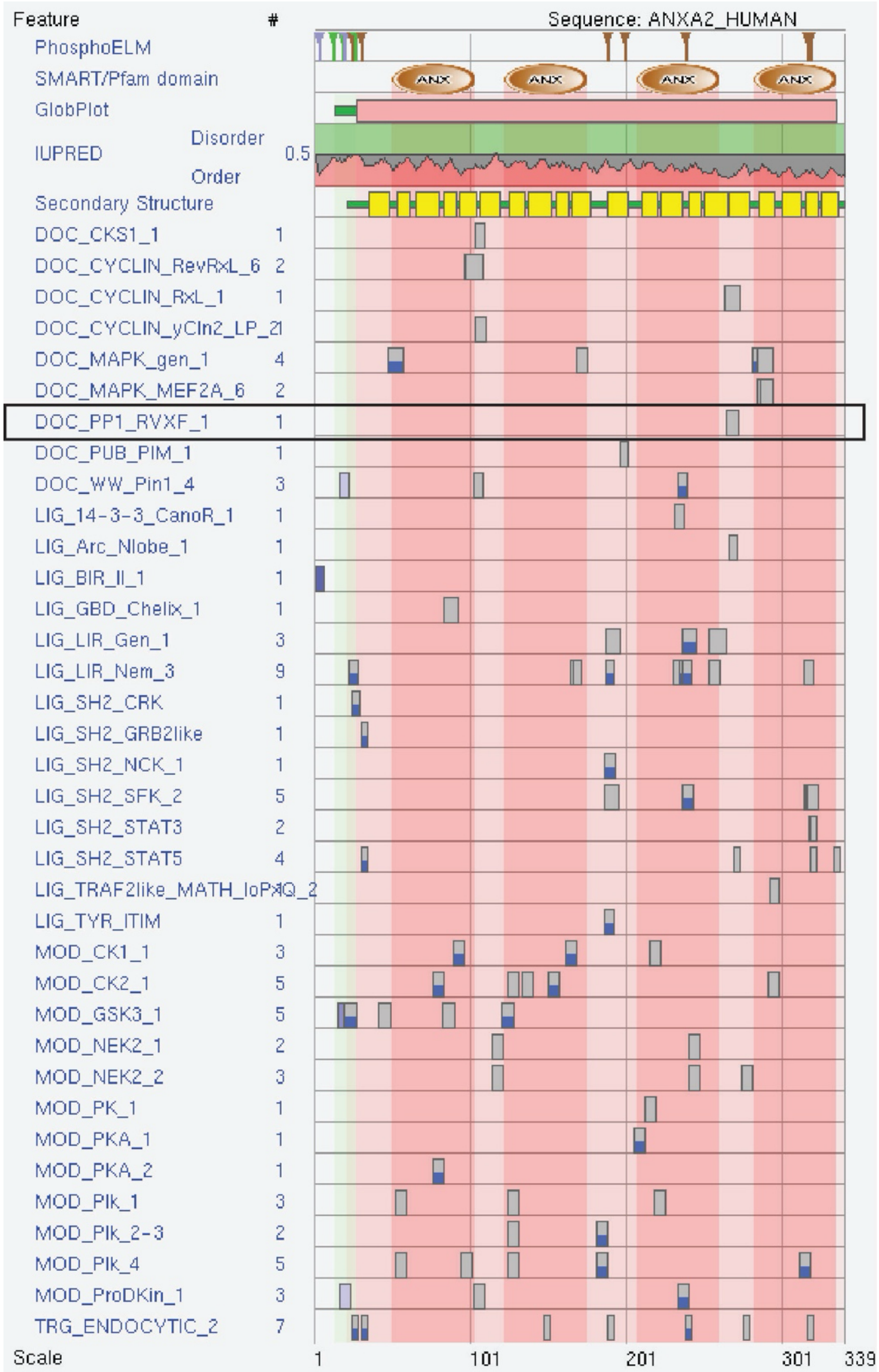


**Supplementary Figure 1: Analysis of putative protein binding domains of ANXA2 using Eukaryotic Linear Motif (ELA) Prediction tool.** Image shown is output from functional site prediction query of ANXA2-HUMAN, restricted to the cytosol compartment. The predicted PP1 binding site is highlighted with a black box.

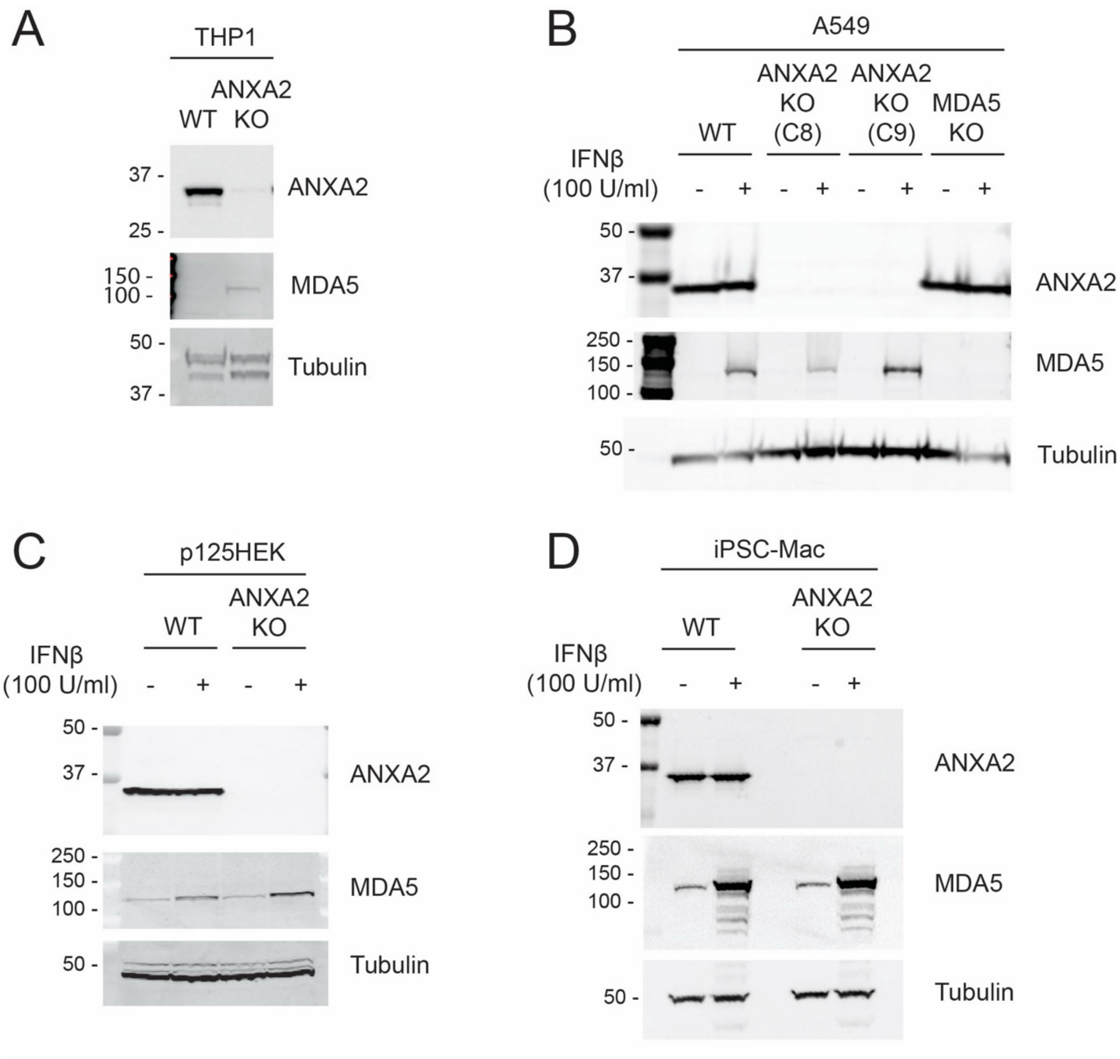


**Supplementary Figure 2: Generation of ANXA2 knockout (KO) cell lines.**
Cell lysates from (A) wild-type (WT) and ANXA2-KO THP1 were analysed by SDS-PAGE and western blot for ANXA2, MDA5 and tubulin (loading control).
(B) WT, ANXA2-KO and MDA5-KO A549 cells, or WT and ANXA2-KO (C) p125HEK cells and (D) iPSC-derived macrophage were untreated or treated with IFNβ (100 U/ml) for 24 hours. Cell lysates were analysed by SDS-PAGE and western blot for ANXA2, MDA5 and tubulin (loading control).

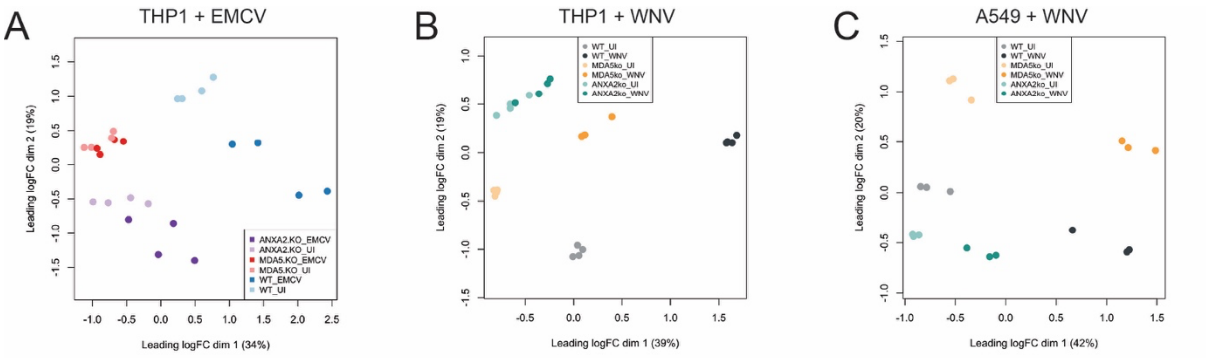


**Supplementary Figure 3: Multidimensional scaling (MDS) plots of transcriptomic data.** Related to (A) Fig 4, (B) Fig 5 and (C) Fig S5, after filtering and normalisation. Individual biological replicates are shown, and colour coded based on genotype (WT, ANXA2-KO or MDA5-KO) and treatment (uninfected (UI), EMCV or WNV). Cell types and treatments are denoted on labels above plots.

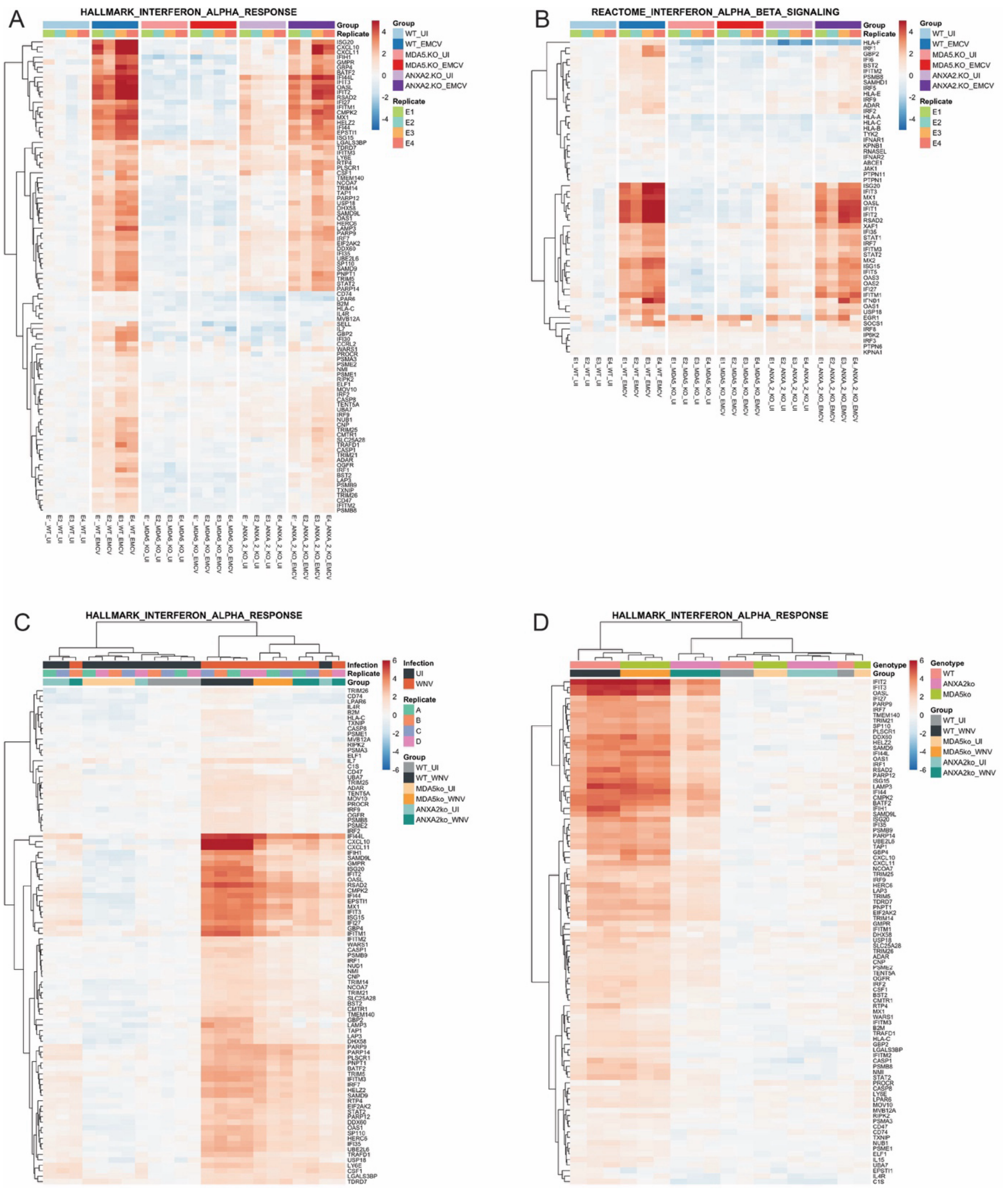


**Supplementary Figure 4. Interferon pathway analysis.**

Related to Figures 4, 5 and S5. Heat maps show the relative expression of all genes in specific gene sets: (A) Hallmark IFN alpha response and (B) Reactome IFN beta response gene sets for THP1 cells infected with EMCV (relative to uninfected (UI)); Hallmark IFN alpha response gene sets for (C) THP1 cells and (D) A549 cells infected with WNV (relative to uninfected).

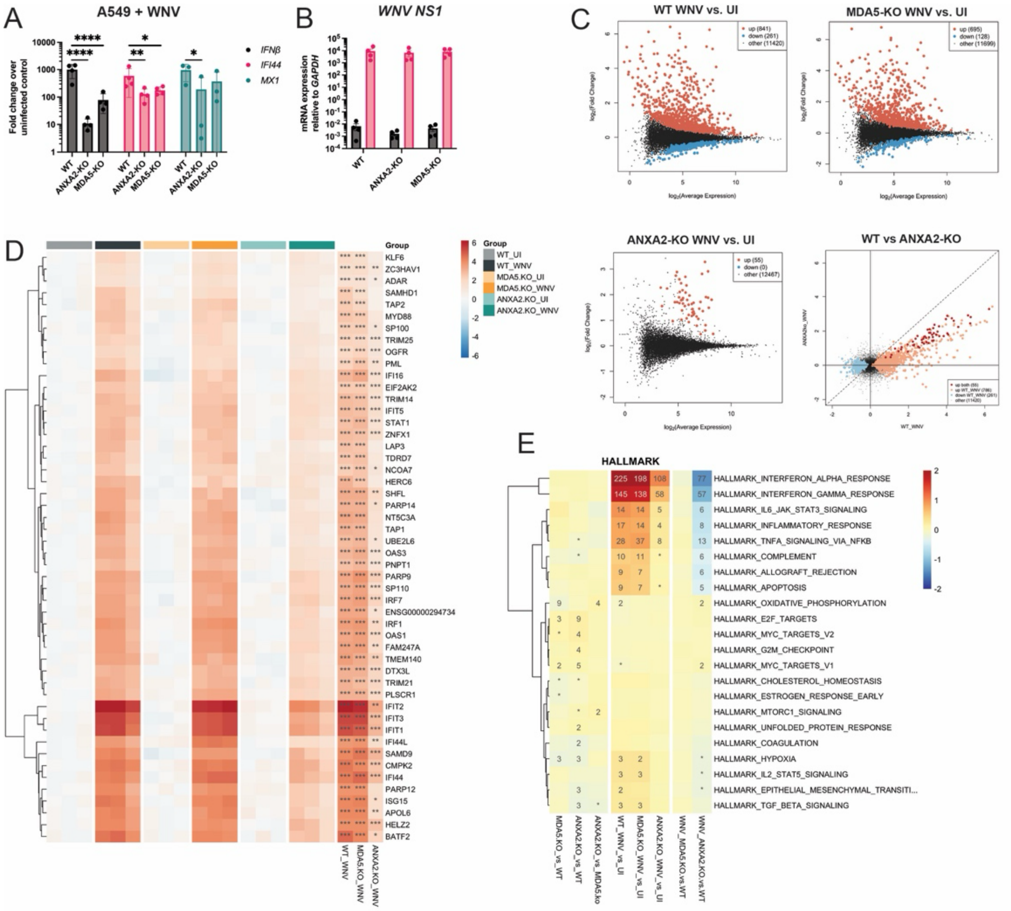


**Supplementary Figure 5: ANXA2 is required for the T1-IFN response to WNV in A549 cells.**

Wild-type, ANXA2-KO and MDA5-KO A549 cells were infected with WNV (MOI = 1) for 24 hours. RNA was extracted and RT-qPCR performed for (A) indicated mRNAs and (B) WNV NS1 RNA, or (C-F) analysed by BRB-seq. Data in (A-B) are shown relative to *GAPDH*, and data in (A) represents fold change over uninfected. Bars in (A-B) indicating the mean values from three independent biological repeats. Statistical significance was determined by 2-way ANOVA.

(C) Differential expression is depicted as $\log_2$ fold changes of all genes between infected (WNV) and uninfected (UI) samples for each genotype versus average gene expression or between ANXA2-KO and WT infected cells directly.

(D) Heat map of relative gene expression of all samples compared to WT uninfected for genes changed in response to infection. Average $\log_2$ fold changes between uninfected and infected samples are indicated in right three columns. Asterisks indicate genes significantly different between comparisons. (* $p < 0.05$, ** $p < 0.01$ and *** $p < 0.001$).

(E) Gene set test results for Hallmark gene set collections. From left to right: columns 1-3 compare between genotypes in uninfected cells, columns 4-6 compare between infected and uninfected cells within each genotype, and columns 7-8 compare between genotypes in infected cells. Colours indicate the average $\log_2$ fold changes of all genes in the gene set (scale truncated to ±2). The significance of each gene set is indicated by the text as the $-\log_{10}$ FDR-adjusted p-value threshold (* denotes adjusted $p < 0.05$, 2 denotes $p < 0.01$, etc.).

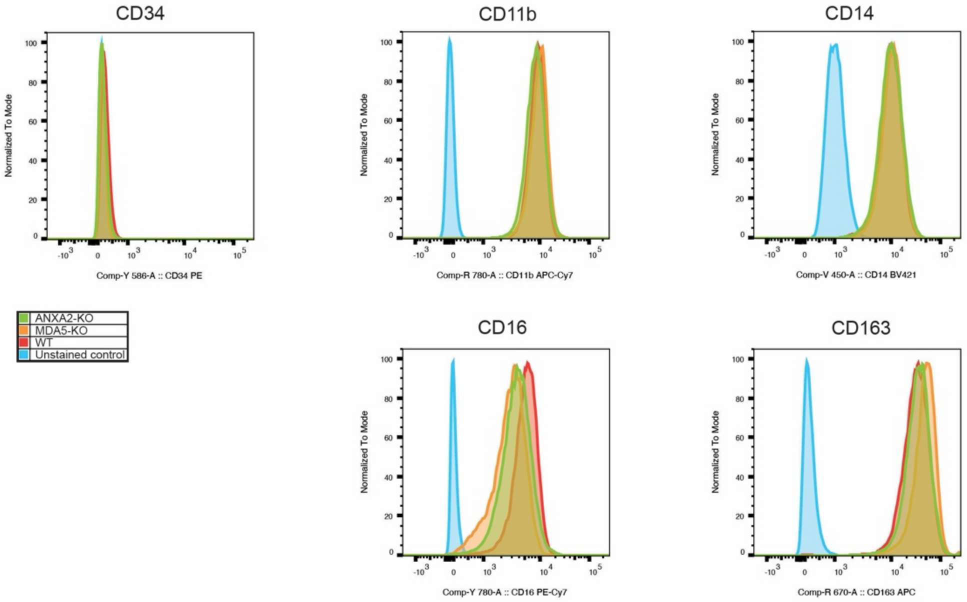


**Supplementary Figure 6. Surface marker staining of iPSC-derived macrophages.** iPSC-derived WT, ANXA2-KO, and MDA5-KO macrophages were stained for hematopoietic stem cell marker CD34, and macrophage markers CD116, CD14, CD16 and CD163. Unstained WT cells were included as a negative control.

| Gene target | Forward primer sequence | Reverse primer sequence |
|---|---|---|
| *CXCL10* | TTCCTGCAAGCCAATTTTGT | TTCTTGATGGCCTTCGATTC |
| *EMCV 5NTR* | GTCTGTAGCGACCCTTTG | CCTTGTTGAATACGCTTGAG |
| *GAPDH* | GGCCATCCACAGTCTTCTG | TCATCAGCAATGCCTCCTG |
| *IAV M1* | AAGACCAATCTTGTCACCTCTGA | TCCTCGCTCACTGGGCA |
| *IFNB* | TGCTCTGGCACAACAGGTAG | CAGGAGAGCAATTTGGAGGA |
| *IFI44* | GTGAGGTCCAAGCTAGAGGAAG | TGCAGCCCATAGCATTCGTC |
| *IL6* | CTCCAGGAGCCCAGCTATGA | CCCAGGGAGAAGGCAACTG |
| *ISG15* | GCGAACTCATCTTTGCCAGT | AGCATCTTCACCGTCAGGTC |
| *MX1* | GGTGGTGGTCCCCAGTAATG | ACCACGTCCACAACCTTGTCT |
| *OAS2* | GAAGCCCTACGAAGAATGTCAGA | TCGGAGTTGCCTCTTAAGACTGT |
| *TNF* | CTCTTCTGCCTGCTGCACTT | TGGGCTACAGGCTTGTCACT |
| *WNV NS1* | TCAAGAATAACTTGGCGATCCA | TCACCTAGGACCGCCCTTT |
| *PAX6* | GCCCTCACAAACACCTACAG | TCATAACTCCGCCCATTCAC |
| *NESTIN* | TGCGGGCTACTGAAAAGTTC | GGCTGAGGGACATCTTGAG |
| *TUBB3* | TTTGGACATCTCTTCAGGCC | TTTCACACTCCTTCCGCAC |

**Table S1.**
Forward and Reverse primer sequences for specific gene ratget detection by qPCR.

**Data S1. (separate file)**
Complete data file for SILAC mass spectrometry, related to Figure 1.
